# Uncovering novel phage defense systems through genomic island analysis in *Streptococcus thermophilus*

**DOI:** 10.64898/2026.10.04.756616

**Authors:** Audrey Leprince, Vincent Somerville, Justine Lefrançois, Geneviève M. Rousseau, Anne M. Millen, Damian Magill, Philippe Horvath, Dennis A. Romero, Sylvain Moineau

## Abstract

Bacteria encode diverse antiviral defense systems, yet the evolutionary reservoirs that generate this immune innovation remain poorly defined. Although genomic context has been instrumental in discovering defense systems, genomic islands (GIs), variable chromosomal regions enriched in defense genes and proteins of unknown function, have not been systematically explored as sources of new bacterial immunity. Here, using comparative genomics of *Streptococcus thermophilus,* we identified 30 defense-associated GIs displaying extensive accessory gene diversity. Investigation of candidate loci from three GIs uncovered 15 previously uncharacterized antiviral systems, including 10 with novel architectures or features, revealing the potential within even a small subset of GIs. Several systems were not located near known defense genes, highlighting that GI-based discovery complements the traditional guilt-by-association approach, revealing systems that would otherwise remain undetected. Together, our findings establish GIs as reservoirs of antiviral innovation and provide a framework for uncovering the diversity of bacterial immune systems.

## Introduction

Bacteria face numerous challenges, among which phage infection represents a major threat. Long-term co-existence with diverse phage communities has driven the evolution of an arsenal of antiviral defense mechanisms^1^. The earliest line of defense acts extracellularly by preventing phage adsorption, typically through receptor modification or masking, thereby blocking infection before the phage genome enters the cell^2^. In addition, bacteria also rely on intracellular defense systems, encoded by individual genes or operons, that disrupt phage replication at various steps of the infection cycle^3^. Over the past years, many such systems have been uncovered, and investigations into their molecular mechanisms have revealed a remarkable diversity of antiviral strategies^4–6^. Although more than 250 defense systems have been described to date, it is widely anticipated that many more remain to be discovered^7,8^.

Although most recently discovered defense systems have originated from studies on Gram-negative bacteria, with only a subset validated in Gram-positive bacteria, the central role of Gram-positive lactic acid bacteria (LAB) in advancing research on anti-phage defense systems is sometimes overlooked. In the food industry, lytic phages represent a persistent threat by infecting LAB used in fermentation processes^9,10^. This continuous selective phage pressure has created a unique opportunity to investigate phage-bacterium interactions at an unprecedented scale and has driven the discovery and characterization of diverse defense mechanisms that protect bacteria from phages circulating in these environments^11,12^. For example, the discovery of numerous abortive infection (Abi) systems originated from studies of the cheese-producing LAB *Lactococcus lactis* and *Lactococcus cremoris*, whose plasmids are particularly enriched in defense systems^13^. Likewise, *Streptococcus thermophilus*, another key dairy starter culture used in yogurt and cheese production, has been instrumental in elucidating the functions and mechanisms of chromosomally-encoded CRISPR-Cas systems^14^. More recently, a few studies have revealed that LAB harbor a great diversity of defense systems shaped by their long-standing evolutionary arms race with their evolving phages^15–17^.

Defense systems have been uncovered through multiple complementary approaches, and examining their genomic localization has proven particularly useful for discovering new systems. A significant advance was the realization that defense systems frequently cluster within bacterial genomes, forming regions now known as defense islands, where multiple defense systems are found within a few kilobases^18^. Exploring genes located near known defense systems within these islands has enabled the identification of many recently characterized defense mechanisms, an approach referred to as guilt-by-association^19,20^. Similarly, mobile genetic elements (MGEs) often contain specific hotspots enriched in defense systems, providing another strategy for narrowing down candidate defense genes^21,22^. These regions, termed defense hotspots, are not defined by the presence of multiple defense systems but rather by their genomic context: they correspond to regions of MGEs flanked by conserved genes, within which at least one defense system is present. The type and number of systems within these regions can vary among closely related MGEs, and exploring defense hotspots containing only uncharacterized genes has led to the identification of several novel defense systems.

At a broader scale, studies in several bacterial species have revealed the presence of chromosomal defense hotspots that relate to genomic islands (GIs), which correspond to variable genomic regions flanked by conserved genes and enriched in defense systems but also in genes coding for proteins of unknown function ^23–25^. These regions are therefore likely to also encode other adaptive functions in addition to antiphage defense, such as drug resistance, pathogenicity traits, and metabolic pathways, thereby contributing to bacterial evolution and ecological adaptation^26,27^. GIs are acquired through horizontal gene transfer (HGT) and are often associated with MGEs and their preferred integration sites (e.g., tRNA loci). The analysis of such GIs enabled the discovery of the Shango defense system in a core defense hotspot of *Pseudomonas aeruginosa*^24^. However, these regions have not yet been systematically explored as reservoirs for novel defense systems.

In this study, we leveraged the chromosomal localization of known defense systems in *S. thermophilus* to uncover 30 GIs associated with phage defense systems, including several hotspots enriched in diverse antiviral systems. Functional screening of candidate genes from three of these GIs, revealed 15 previously unknown defense systems that provide protection against *S. thermophilus* phages, the majority of which have not been characterized in any bacterial species. These findings establish GIs as important reservoirs of bacterial immunities.

## 1. Results

### *S. thermophilus* defense systems localize in 30 genomic islands

We previously analyzed the defensome of *S. thermophilus*, identifying 28 defense systems primarily encoded on the chromosome^17^. CRISPR–Cas and RM systems were the most abundant defense systems. Type II-A CRISPR–Cas systems were identified in 100% of the strains, while type I RM systems were present in nearly 95% of the strains. Both were therefore considered core defense systems. Most of the remaining systems were classified as accessory, being present only in a subset of strains (**Supplementary Figure S1**). In this study, we focused on 81 complete *S. thermophilus* genomes, harboring 23 known and 12 candidate defense systems, to determine the chromosomal locations of known defense systems and identify GIs enriched in defense systems. To this end, we first performed a pan-genome analysis, which allowed us to define single-copy non-defense core genes (i.e., present in 100% of the strains) (**Figure 1A**). Each validated defense system, as well as candidate systems predicted by PADLOC^28^, was then analyzed across the 81 genomes to identify its nearest upstream and downstream single-copy non-defense core genes. This approach enabled the delineation of GIs composed of accessory genes, including defense genes. Finally, all core non-defense gene pairs flanking GIs were mapped across the 81 strains, allowing further characterization of GIs associated with known anti-phage defense.

**Figure 1.**
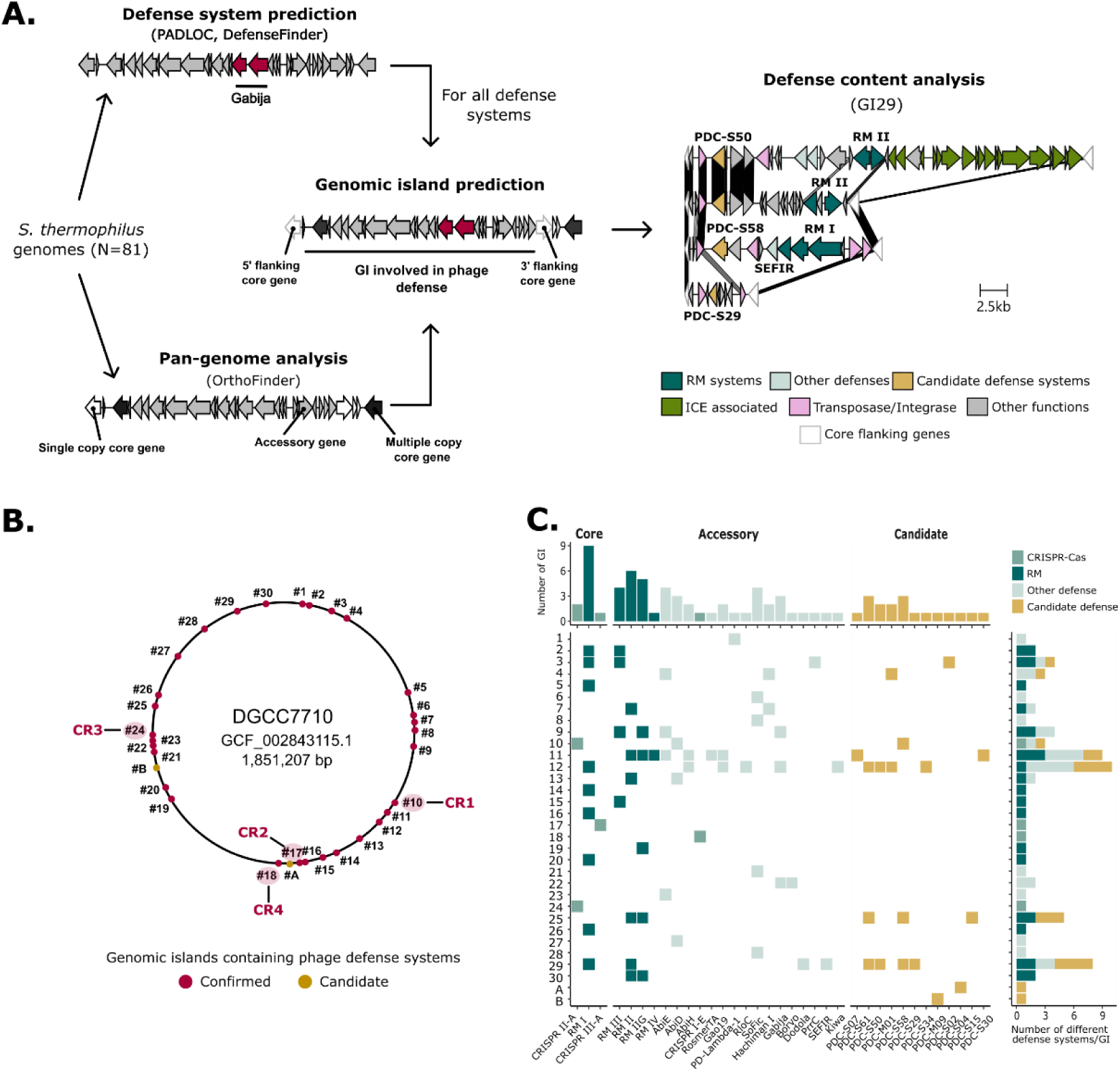
Delimitation of genomic islands (GIs) associated with phage defense systems in *S. thermophilus*. **A.** Workflow illustrating the strategy used to identify GIs containing phage defense systems across 81 complete *S. thermophilus* genomes. Core host genes were defined as those present in all analyzed genomes, and single-copy core genes served as anchors to delineate GI boundaries. **B.** Localization of the identified GIs in the reference genome *S. thermophilus* strain DGCC7710. Thirty GIs (1–30) were defined based on the presence of known (previously described) defense systems (red). Two additional regions containing only candidate defense systems predicted by PADLOC were designated as candidate GIs (yellow, A and B). GIs linked to the four CRISPR-Cas systems in *S. thermophilus* (CR1 and CR3: type II-A; CR2: type III-A; CR4: type I-E) are highlighted. **C.** Defense system occupancy across the 32 GIs identified from 81 complete *S. thermophilus* genomes. The core known defense systems (CR1/CR3, CR2, and type I RM) were defined as those present in >70% of genomes, whereas accessory systems occurred in <70%. Candidate defense systems correspond to PADLOC predictions. The upper bar graph shows the number of GIs in which each defense system is found, and the right bar graph indicates the number of distinct defense systems per GI.

We identified 30 GIs (numbered 1-30) containing genes coding for anti-phage mechanisms, distributed throughout the chromosome of all 81 genomes (**Figure 1B**). Contrary to what was observed in *Escherichia coli* and *P. aeruginosa*, none of the GIs were located within tRNA loci (**Supplementary Data S1**)^23,24^. While some defense systems (11/23) were restricted to a single GI, others (12/23) were present in multiple GIs (**Figure 1C**). Notably, RM systems were particularly widespread with type I RM systems distributed across nine GIs. On the contrary, the four CRISPR arrays were always located in the same GI (**Figure 1B-C**). While most GIs contained only one (17/30) or two (5/30) types of known defense systems, several, including GI9, GI11, GI12, and GI29, harbored a more diverse repertoire with at least four distinct known defense systems. PADLOC candidate systems, initially identified because they were embedded within known defense systems^29^, were primarily found within the defined GIs, therefore occupying the same locations as validated defense systems. This pattern further supports their potential defensive role. Of the 12 PADLOC candidate systems identified in *S. thermophilus*, only two (PDC-M09 and PDC-S04) were not located in the GIs, thus defining two candidate GIs (A and B) potentially involved in phage defense (**Figure 1B-C**).

### Defense associated GIs exhibit diverse gene content

Beyond their diversity in defense systems (**Figure 1C**), the GIs of *S. thermophilus* associated with phage defense also varied in size and content. In this study, a GI was defined solely by the presence of at least one defense system (validated or candidate) at the corresponding locus in at least one strain. As a result, three distinct occupancy states were observed across strains (**Figure 2A**). First, 30% of the GIs (774 out of 2,592 GIs in our 81-strain dataset) were empty, with the two flanking genes encoded directly adjacent to one another and no gene between them. Second, 48% (1251/2,592) contained only genes unrelated to known defense systems (i.e., occupied by 0 defense system). Third, 22% of GIs (567/2,592) harbor genes that include at least one defense system (i.e., occupied by 1, 2, or 3 defense systems). Additionally, we looked at the variation in GI size and gene content focusing on the functional category and gene mobilisation (**Figure 2B-D**). Overall, genes located within GIs represent 24–50% of the accessory genome (genes not shared by all strains) (**Figure 2C**).

**Figure 2.**
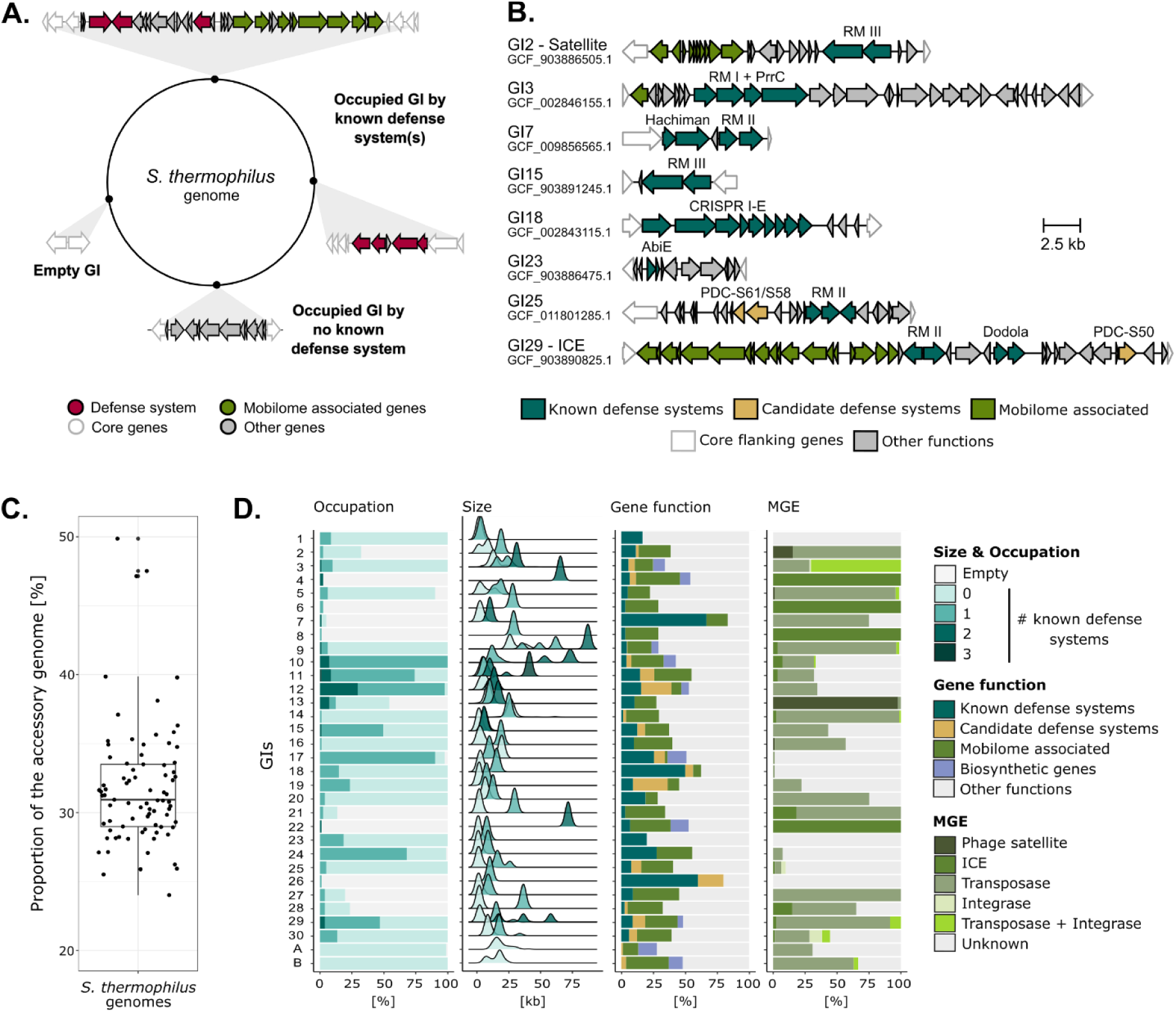
Gene content of GIs associated with phage defense in *S. thermophilus*. **A.** Schematic representation of possible occupation status for the defense system associated with GIs **. B.** Representative genetic organization of GIs occupied by known defense systems. GIs corresponding to phage satellite insertions or integrative and conjugative elements (ICEs) are indicated. Strain accession numbers are provided below each GI. **C.** Contribution of GI genes to the accessory genome across 81 complete *S. thermophilus* genomes. Each dot represents the proportion of accessory genes located within GIs for a single genome. **D.** Characteristics of GIs identified in the 81 genomes analyzed. The bar graph and density plot depicting GI occupancy and size are colored according to GI occupation: “empty” (no genes between flanking core genes), “0” (genes present but no known defense systems), or numbers (1–3) corresponding to the number of defense systems detected. Additional panels summarize the nature of mobile genetic elements (MGEs) or mobilome genes (standalone transposase and/or integrase) associated with each GI and the functional categories of genes within GIs. In these cases, the percentages indicated in relation with GI that are occupied (presence of at least one gene).

While six GIs (GI4, GI6, GI7, GI8, GI22, and GI26) were occupied (i.e., presence of at least one gene in the GI) in only a few closely related strains, most GIs were occupied in at least 75% of the strains and predominantly contained genes unrelated to known defense systems (**Figure 2D** and **Supplementary Figure S2**). GI10 consistently harbored at least one defense system as it is the location of the conserved CRISPR1 locus (type II-A CRISPR-Cas system), which is a core defense system in *S. thermophilus*^17^. The neighboring GI11 and GI12 also encoded known defense systems in more than 70% of the strains. Overall, these GIs are relatively small but are primarily composed of validated and PADLOC candidate defense systems, with few or no genes involved in biosynthetic functions, which suggests that they are mainly defense islands (**Figure 2D**). Together, GI10, GI11, and GI12 form a larger region of approximately 50 kb that is particularly rich in defense systems (**Supplementary Figure S3A**). GI17 and GI24 display similar patterns, as they are also frequently occupied by defense systems.

GI sizes ranged from a few kilobases to insertions of up to 80 kb, the latter typically corresponding to large integrated Integrative and Conjugative Elements (ICEs) (**Figure 2D**). While some GI were specific to a single type of insertion (i.e., identical or near-identical gene content across strains), most of them exhibited a diverse array of insertions whose gene content varied between strains (**Supplementary Figure S3B-C**). Comparative analyses further revealed that several GIs were formed through multiple, discrete integration events rather than a single insertion (**Supplementary Figure S3C**). In addition to genes involved in defense and mobility, several GIs also carried biosynthetic gene clusters as well as genes of various functions, highlighting their functional heterogeneity (**Figure 2D**).

To investigate the potential mechanisms driving GI gene diversity and mobilization, we analyzed MGEs and markers of HGT associated with these regions (**Figure 2D**). *S. thermophilus* is known to be enriched in transposases, which contribute to the species domestication^30^. Consistently, transposases were the most abundant HGT marker, occurring in 24 GIs. ICEs also emerged as major contributors to GI mobilization and phage defense. Although only one RM system encoded within *S. thermophilus* ICEs has been functionally characterized^31^, *S. thermophilus* ICEs encode a wide diversity of defense systems (**Supplementary Figure S4A**) and were found in 12 GIs. Five of these (GI4, GI6, GI8, GI21, and GI22) correspond to specific ICE integration sites. Phage satellites, which have been associated with phage defense in *S. thermophilus*^32^ (**Supplementary Figure S4B**), were integrated within GI2, GI5, GI13, and GI16; notably, GI13 appears to be a major integration hotspot. Finally, for 47.3% GI-strain combinations (860 out of 1,818 GIs occupied by genes, either including known defense systems or not), no clear mobilization mechanism could be identified.

### *S. thermophilus* GIs can be leveraged to uncover novel defense systems

We next aimed to identify novel defense systems within *S. thermophilus* GIs. Many defense systems have been discovered through a guilt-by-association approach. Additionally, systematic exploration of hotspots in MGEs, including those lacking previously characterized defense systems, has proven effective for uncovering new mechanisms^21^. In this work, the *S. thermophilus* GIs, as we have defined them, were investigated for the presence of uncharacterized defense systems including some that are not found in the vicinity of known defense systems.

Three GIs (GI3, G11, and GI12) were selected to investigate new anti-phage systems for experimental validation against members of the five viral genera known to infect *S. thermophilus*^33^ (**Figure 3A**). Candidate genes or operons were selected manually, excluding predicted pseudogenes and genes with involved in biosynthetic clusters and HGT and prioritizing those of unknown function. Four candidate systems from GI3 were chosen exclusively from strains in which this GI contains no known defense systems and thus could not have been selected through a guilt-by-association approach. Additionally, 14 candidate systems were chosen from GI11 and GI12 as these GIs showed diverse contents of known defense systems and were consistently occupied across all strains (**Figures 1C** and **2D**). Except for two, the candidate systems selected within these two GIs co-localized with known defense systems (N = 12). Notably, candidates from GI12 were predominantly (9/10) embedded within the subunits of a type I RM system, an arrangement previously reported for other defense systems and forming the basis of candidate prediction in PADLOC^29^. In total, 18 candidate systems were selected for experimental validation (**Supplementary Data S2**).

**Figure 3.**
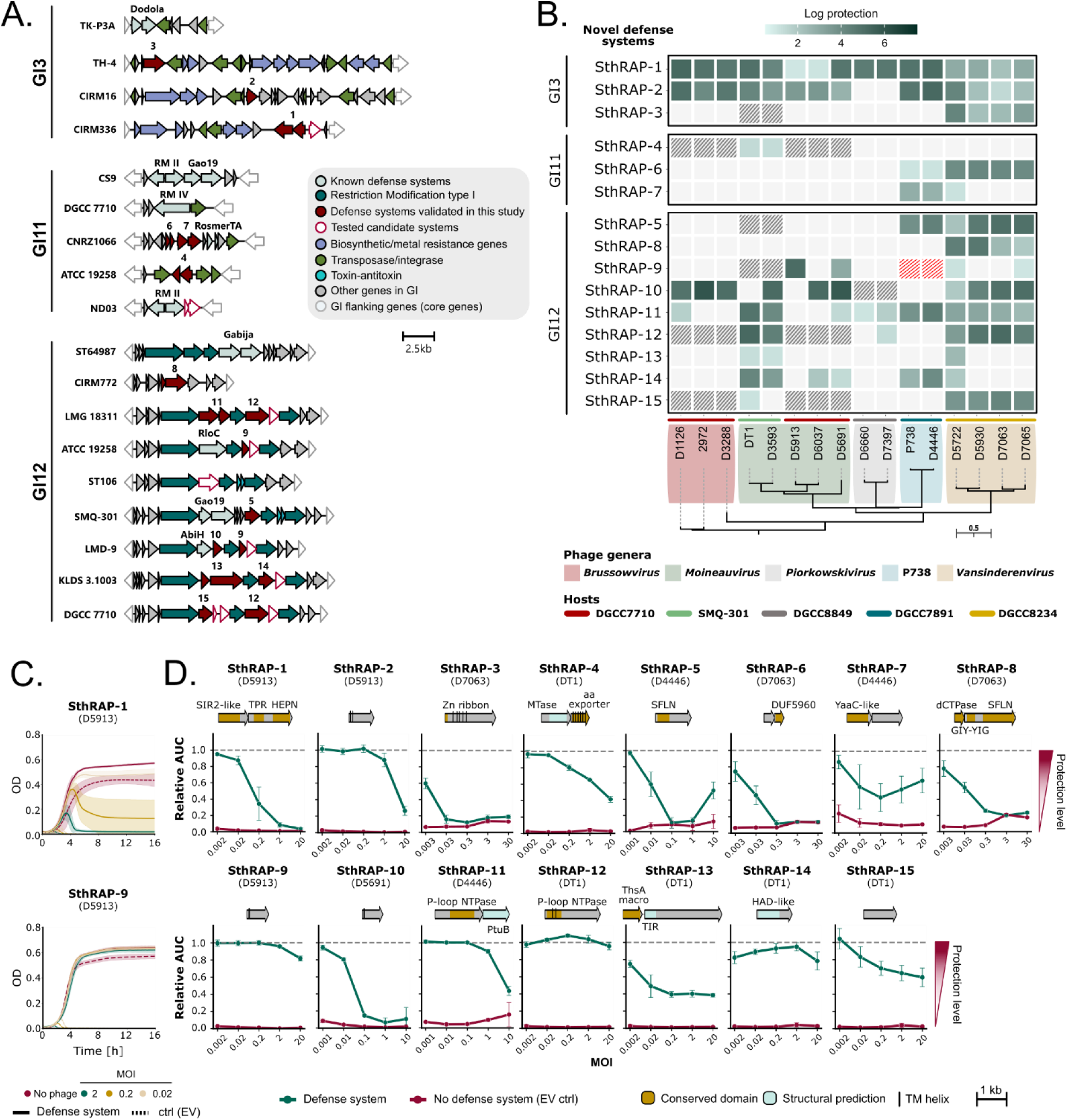
Defense activity of SthRAP systems in *S. thermophilus*. **A.** Genetic organization of the genomic islands (GIs) analyzed in this study to identify novel defense systems. Numbers indicated above some genes correspond to the newly confirmed SthRAP (*<u>S</u>treptococcus* <u>th</u>ermophilus <u>R</u>esistance <u>A</u>gainst <u>P</u>hage) presented in the other panels. **B.** Heatmap summarizing anti-phage activity of validated defense systems against virulent *S. thermophilus* phages. Activity was assessed on solid media by spot test assays, with a minimum threshold of ≥1-log reduction in phage titer. Data represent three biological replicates. Striped cells indicate conditions not tested because the defense system was endogenous to the strain (grey) or transformation was unsuccessful (red). **C.** Representative phage infection assays in liquid media showing anti-phage activity over 16 hours at 37°C for three multiplicities of infection (MOIs) of phage D5913 in strain DGCC7710 expressing either SthRAP-1 or SthRAP-9. Control (ctrl) corresponds to *S. thermophilus* strain DGCC7710 carrying the empty vector (EV) pTRKL2. Curves represent the mean value of three biological replicates, with shaded areas indicating confidence intervals. **D.** Summary of phage infection assays in liquid media for various SthRAP systems at five MOIs. Defense activity is expressed as the ratio between the area under the curve (AUC) of infected cultures and the non-infected control cultures. A relative AUC of 1 indicates full protection (growth comparable to the absence of phages), whereas 0 indicates no protection (complete growth inhibition). The tested phage is indicated in brackets and the relative AUC for the control strain carrying the empty vector (EV) is shown for comparison (red). Data represent the mean of three biological replicates, with error bars showing standard deviation. Genetic organization of SthRAP systems is displayed above each plot, with conserved domains (InterProScan) and structural homologies (DALI) annotated. Abbreviations: aa, Amino acid; dCTPase, dCTP pyrophosphatase; DUF, Domain of unknown function; GIY-YIG, GIY-YIG endonuclease; HAD, Haloacid dehalogenase; HEPN, Higher eukaryotes and prokaryotes nucleotide-binding; MTase, Methyltransferase; NTPase, Nucleoside-triphosphatase; PtuB, Septu subunit B; SFLN, Schlafen; SIR2, Sirtuin; ThsA, Thoeris subunit A; TIR, Toll/interleukin-1 receptor/resistance protein; TM, Transmembrane; TPR, Tetratricopeptide.

Selected candidates were cloned onto a low copy plasmid and tested against phages as previously described^17^. Overall, 15 candidates out of the 18 displayed defense activity against at least one tested phage, providing protection ranging from 1 to 6 log units in plate assays (**Figure 3B** and **Supplementary Figure S5**). These newly discovered defense systems were renamed SthRAP for *<u>S</u>treptococcus <u>th</u>ermophilus* <u>R</u>esistance <u>A</u>gainst <u>P</u>hages. Similar to known defense systems^17^, SthRAP systems exhibited variable specificity. Broad spectrum systems such as SthRAP-1 and SthRAP-2 conferred protection against all tested phages from five and four genera, respectively. In contrast, most other SthRAP systems displayed narrower activity, targeting phages from only one genus (SthRAP-3, SthRAP-4. And SthRAP-9), two genera (e.g., SthRAP-5 and SthRAP-6), or even only a few individual phages (e.g., SthRAP-9). Most candidates (5 out of 6) originating from GIs lacking known defense systems were validated as functional defense systems, specifically SthRAP-1, SthRAP-2, SthRAP-3, SthRAP-4, and SthRAP-8. This supports the idea that such GIs are promising locations for the discovery of uncharacterized defense mechanisms. In addition, most candidates located within type I RM loci (8 out of 9) also displayed defense activity, further expanding the diversity of defense systems that can be found embedded within other defense systems ^29^. Interestingly, although 13 of the validated systems received full (10/13) or partial predictions (3/13) as potential defense systems by PADLOC^29^ or DefensePredictor^7^, SthRAP-3, SthRAP-4, and SthRAP-6 were not highlighted by either tool (**Supplementary Data S2**). Conversely, two candidate systems that were predicted to be defensive did not show detectable activity against the phages tested (**Supplementary Data S2**).

The contribution of individual genes to defense activity within multi-gene systems was assessed by testing each gene separately (**Supplementary Figure S6**). Most two-gene systems were confirmed. However, in SthRAP-9 and SthRAP-12, the shared *cdsB* (PDC-S61, also associated with SthRAP-14) was dispensable, and defense depended entirely on *cdsA*. Likewise, the anti-phage activity of SthRAP-15 was solely attributable to *cdsA*. Defense activity was further validated in liquid media across multiple MOIs (**Figure 3C-D**). This approach confirmed the activity of systems that showed only modest protection in plate assays (<3-log), such as SthRAP-4, SthRAP-7, and SthRAP-13. Liquid assays also suggested differences among systems: some provided protection at high MOIs, consistent with direct defense, whereas others were protective only at low MOIs, suggestive of an abortive infection (Abi) phenotype^34^ (**Supplementary Figure S7**).

Altogether, the analysis of three GIs led to the identification of 15 novel defense systems in *S. thermophilus*, further expanding the immune repertoire of this species and establishing GIs as defenses-rich rich genomic regions.

### Some new SthRAP systems have similarities with known defense systems

Functional annotation based on sequence and structural analysis with AlphaFold3 (AF3) was performed for the validated SthRAP systems, which included multimer and ligand binding prediction when relevant (**Figure 4**, **Supplementary Figure S8**, and **Supplementary Data S3**). Five of the 15 systems display structural similarities with defense systems that were previously identified.

**Figure 4.**
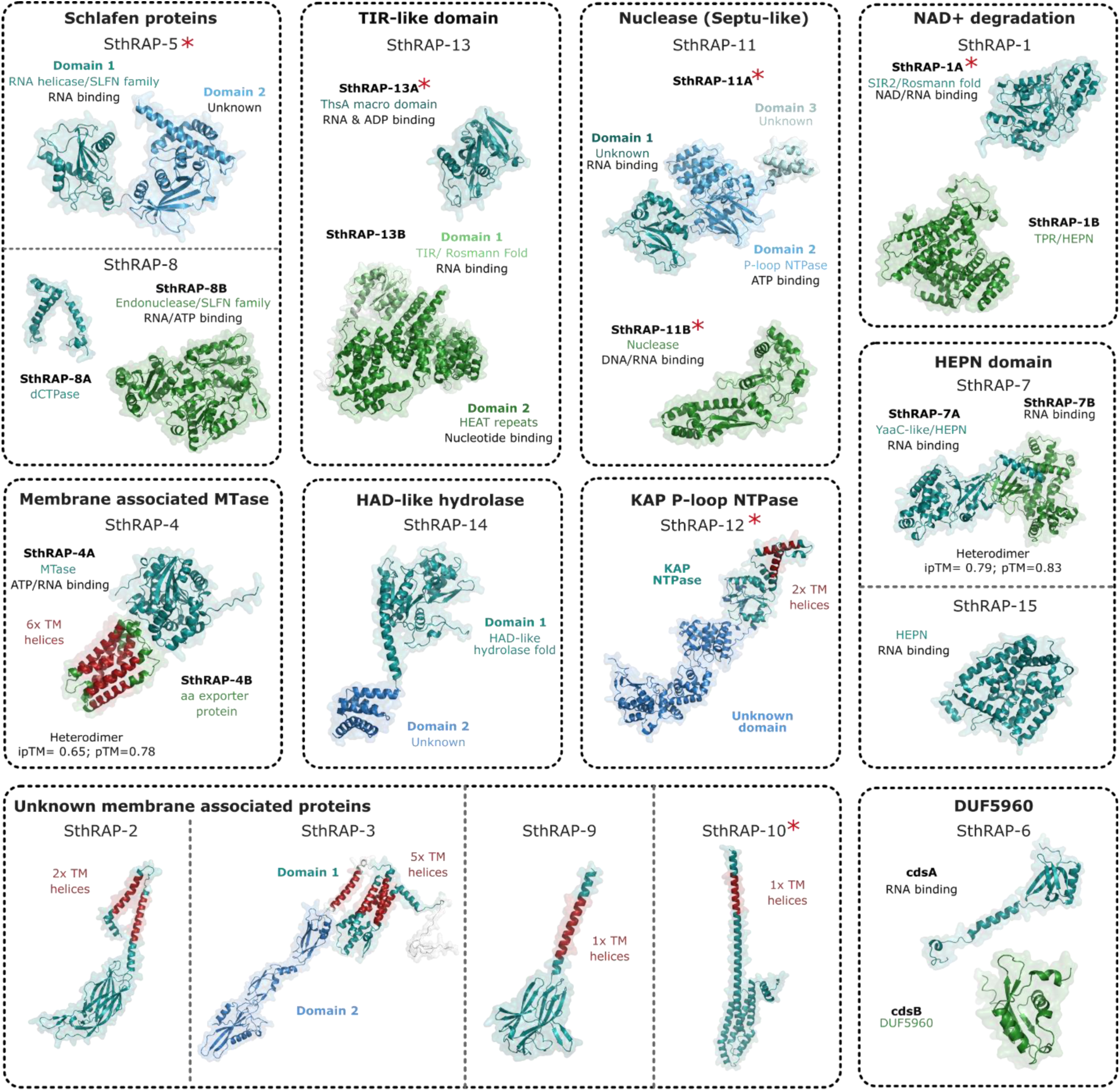
Structural models of SthRAP systems predicted by AlphaFold3. Predicted protein structures for each SthRAP system are shown, with transmembrane (TM) helices highlighted in red and component proteins in blue and green, respectively for a multi-protein system. Distinct color shading denotes predicted domains. Conserved folds and domains implicated in putative mechanisms of action, as well as predicted interactions with nucleic acids or ligands, are annotated. Proteins exhibiting structural similarity to characterized defense systems or defense effectors are indicated with a red star (see **Supplementary Fig. SG**). Additional details on functional and structural analyses are provided in **Supplementary Data S3** and **Supplementary Fig. S8**.

Two SthRAP systems (SthRAP-5 and SthRAP-12) are homologs of defense systems found in *Enterococcus*^35^. SthRAP-5 contains a N-terminal Schlafen (SLFN) domain, and structural superimposition with the enterococcal SLFN system indicates that they very likely represent the same system (RMSD 6.7 over 256 out of 404 aa) (**Supplementary Figure S9A**). SLFN proteins also mediate antiviral responses in eukaryotes^36–38^ and were recently shown to protect bacteria from phages by tRNA cleavage^39^. In SthRAP-5, the SLFN domain likely interacts with RNA (ipTM = 0.83) and is fused to a putative C-terminal sensor domain of unknown function. SthRAP-12 matches an enterococcal KAP NTPase^35^, as AF3 structural comparison reveals near-perfect superimposition (RMSD 3.3 over 576 out of 622 aa), except for two missing N-terminal helices in SthRAP-12 (**Supplementary Figure S9B**). This system comprises a P-loop NTPase domain of the KAP family (IPR011646) characterized by two embedded transmembrane (TM) helices. The C-terminal region lacks detectable homology but has been proposed to function as an effector. In addition to the enterococcal membrane-associated KAP NTPases recently identified^35^, similar domains have been described in the F plasmid-encoded PifA protein^40^ and Tmn^20^, suggesting that they constitute a broad class of defense systems.

SthRAP-11 shows strong similarities with Septu type I (**Supplementary Figure S9C**). An overlap between Septu and PDC-M01, the latter being a PADLOC defense candidate matching SthRAP-11, was already highlighted^29^, and 63% (1133/1796) of the systems identified as SthRAP-11 here also match Septu (**Supplementary Data S4**). In the absence of phage infection, ATP binding to the ATPase PtuA inhibits the nuclease activity of PtuB, maintaining the inactive state^41^. During phage infection, a decrease in intracellular ATP levels likely lifts this inhibition, activating PtuB, which cleaves phage DNA. Subunit SthRAP-11B is structurally homologous to PtuB (**Supplementary Figure SGC**) and predicted to bind RNA (ipTM = 0.92). As for the subunit SthRAP-11A, the central region is an AAA ATPase and displays a fold similar to PtuA (RMSD 5.1 over 168 out of 536 aa). However, SthRAP-11A also includes additional N- and C-terminal domains for which only remote structural similarities (best DALI Z-scores ca. 4) with a few residues across different proteins were detected, making functional predictions uncertain. Unlike Septu, SthRAP-11 is not predicted to form a heterocomplex, but we hypothesize that it functions through a related mechanism, possibly representing a novel subtype of the Septu defense system.

SthRAP-13 shares similarities with type II Thoeris, in which NAD⁺-derived signaling molecules produced by the ThsB TIR domain activate the ThsA effector via its macro domain, inducing membrane permeabilization to block phage replication^42,43^. Subunit SthRAP-13A is structurally homologous to ThsA macro domain (RMSD = 2.5 over 192 out of 214 aa) involved in ADP-ribose binding but does not include a cognate effector domain (**Supplementary Figure S9D**). Component SthRAP-13B contains an N-terminal domain that adopts a Rossmann fold and is structurally homologous to TIR domains (**Supplementary Figure S9D**). As for the C-terminal domain, it consists of a succession of α-helices with homologies to diverse eukaryotic proteins harboring HEAT repeats, which are typically involved in protein-protein interactions or DNA binding^44,45^.

Finally, SthRAP-10 is a TM protein predicted to form a hexamer, which exhibits structural homology to VP1848 (RMSD 8.6 over 200 out of 235 aa), a defense system identified in *Vibrio parahaemolyticus*^46^ (**Supplementary Figure S9E**). Despite these similarities, they occur in distinct bacterial phyla: SthRAP-10 is restricted to the *Bacillota*, whereas VP1848 is found in *Pseudomonadota* (**Supplementary Figure S10A**). SthRAP-10 features a C-terminal globular domain that may contribute to signal transduction, however there are few insights into its putative mechanism of action.

### Most SthRAP systems represent previously uncharacterized systems

In addition to those exhibiting similarities to previously characterized systems, ten SthRAP systems appear to be new defense systems. They display unique structural and/or functional features, suggesting their mechanisms of action differ from those described to date.

SthRAP-8 is composed of a dCTPase (SthRAP-8A), suggesting an involvement of dNTP hydrolysis similar to previously characterized dCTP deaminase– and dGTPase-based defense systems^47^. The second protein (SthRAP-8B) harbors a N-terminal GIY-YIG endonuclease fused to a SLFN domain, predicted to interact with RNA (ipTM = 0.88–0.91). Although GIY-YIG endonucleases typically function as site-specific nucleases^48^, AF3 did not predict DNA interactions.

SthRAP-1A is characterized by a SIR2 fold, a structural motif commonly found in defense systems that mediate NAD⁺ degradation, indicating a potential Abi mechanism (**Supplementary Figure SGF**). SthRAP-1B is predicted to form a dimer and harbors a tetratricopeptide repeat (TPR) fold associated with a HEPN domain, suggesting a role in interacting with viral proteins and/or RNA, and potentially acting as the sensor protein. Additionally, two other systems have similarities with HEPN domains. In SthRAP-15, the N-terminal RNA interacting (ipTM=0.91) region shows weak structural similarity with the HEPN domain of Csm6 (6TUG-B, Z-score 4.2) and the toxin of HEPN-MNT toxin-antitoxin (TA) system (5YEP-B, Z-score 4.3). In SthRAP-7, the YaaC-like protein (IPR026988, SthRAP-7A) is also structurally related to HEPN-containing proteins (2HSB-A and 5YEP-B, Z-score 7-6), while SthRAP-7B shows similarity with parts of phage tail proteins, mostly tail fiber (Z-scores 8-6, 6u5h-A and 3d37).

SthRAP-14 is a predicted two-domain tetramer. Its N-terminal domain shows strong structural homologies (top hits Z-scores > 15) to members of the HAD-like hydrolase superfamily, which notably includes phosphoserine phosphatases, homoserine kinases, and dehalogenases^49^. It also harbors a small C-terminal domain (95 amino acids) that consists of five α-helices.

The hallmark protein of SthRAP-4 is a S-adenosyl-L-methionine (SAM)-dependent methyltransferase (MTase) (SthRAP-4A, Z-score > 15) whose fold resembles that of tRNA (4GEK) and small-molecule (4QDK) MTases. AF3 predicts interactions with ATP (ipTM=0.81) and RNA (ipTM=0.75), supporting a SAM-dependent MTase involved in tRNA methylation. The second protein, SthRAP-4B, forms a compact bundle of TM helices, which associate with the MTase to form a heterodimer (ipTM= 0.65).

SthRAP-6 consists of two small proteins. SthRAP-6A (121 aa) adopts a fold resembling that of RNA chaperones such as Hfq (3HFO, Z-score 9.2) and its homologue YaeO (6JIE, Z-score 8.6). The second protein, SthRAP-6B (108 aa), contains a DUF5960 domain and exhibits weak structural similarity (Z-scores 3.5–4.4) to toxins from several TA systems, including ribosome-dependent RNases that cleave tRNA (6CP9) or translating mRNA (5MJE, 3G5O)^50^.

Lastly, SthRAP-2, SthRAP-3, and SthRAP-9 are TM proteins that lack strong structural homologies with proteins of known function. SthRAP-2 is predicted to assemble into an octamer with a C-terminal cytoplasmic domain that may mediate protein–protein interactions. SthRAP-3, on the other hand, features an N-terminal zinc-ribbon domain and a C-terminal domain of unknown function. SthRAP-9 assembles into a hexamer (ipTM=0.82) and harbors a C-terminal globular domain with weak structural similarity to C2 domains of Synaptotagmin-1 (5CCH-E, Z-score 6.8) and KIBRA (6FJC-A, Z-score 6.5), which mediate calcium-dependent phospholipid binding in eukaryotes^51,52^. AF3 modelling of SthRAP-9 predicted calcium-binding capability (ipTM = 0.9).

### SthRAP systems are broadly distributed across bacterial phyla

To evaluate the prevalence of SthRAP systems, we built Hidden Markov Models (HMM) profiles and performed sequence similarity searches across complete bacterial genomes (RefSeq June 2025, N= 50,676), ensuring gene co-localization for multi-component systems. Consistent with previous large-scale analyses of the prevalence of defense systems^53^, most SthRAP systems occurred in fewer than 1% of the genomes (**Figure 5A**). Nonetheless, several systems were comparatively more widespread. SthRAP-6 and SthRAP-5 were each detected in 1.2% of genomes, while SthRAP-8 and SthRAP-11 were found in 4.6% and 3.4% of genomes, respectively. Most SthRAP homologs were broadly distributed across diverse bacterial phyla, except for SthRAP-2, SthRAP-10, and SthRAP-13, which were restricted to the *Bacillota* (**Figure 5A**) with the latter two being only found in streptococci.

**Figure 5.**
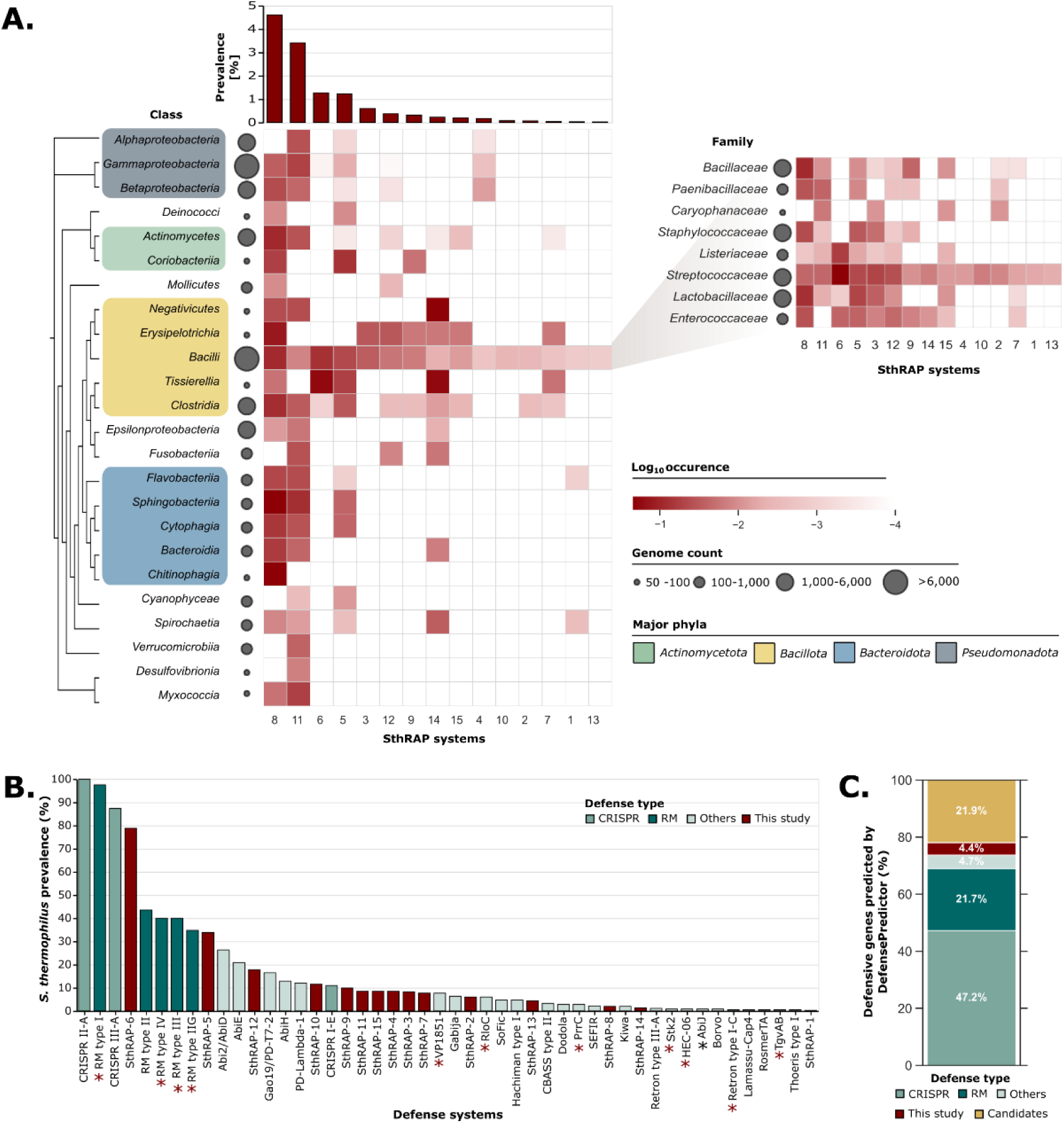
Distribution of SthRAP systems in other bacterial genomes. **A.** Distribution of SthRAP systems in complete bacterial genomes (RefSeq June 2025, N=50,676). The bar graph depicts the overall prevalence of SthRAP systems in the dataset, while heatmaps show their phylogenetic distribution within classes or families (of the *Bacilli*) represented by ≥ 50 genomes. Occurrence in each taxon is normalized to the total number of genomes in that taxon and expressed as log_10_ values. **B.** Prevalence of 48 defense systems across *S. thermophilus* genomes (RefSeq June 2025, N=364). SthRAP systems were identified using hidden Markov models (HMMs) and MacSyFinder, and known systems were predicted with DefenseFinder and PADLOC. Defense systems lacking activity against *S. thermophilus* phages are indicated by a black asterisk and systems not yet experimentally tested in *S. thermophilus* are marked with a red asterisk. **C.** Classification of genes predicted as defensive (DefensePredictor; log-odds > 7.2) in *S. thermophilus* genomes (total of 11,143 genes in 364 genomes), showing their assignment to known systems (CRISPR, RM or others), new systems identified in this study, or uncharacterized candidates corresponding to genes predicted as defensive by DefensePredictor which do not correspond to validated systems.

Next, we performed an in-depth prevalence analysis within *S. thermophilus* by combining the search for SthRAP systems with that of previously characterized defense systems. Using updated prediction tools and a dataset of 364 *S. thermophilus* genomes from RefSeq (June 2025), we generated an updated overview of the *S. thermophilus* defensome. Altogether, integrating SthRAP systems with those identified by prediction tools, revealed 48 defense systems in *S. thermophilus*, substantially expanding the diversity compared to our previous study, which reported 28 systems ^17^ (**Figure 5B**). The average number of defense systems per strain increased from 7.5 to 9.7, with individual strains harboring between 4 and 22 systems (**Supplementary Figure S10B**). SthRAP-6, SthRAP-5, and SthRAP-12 were particularly prevalent, each occurring in more than 15% of strains. Notably, SthRAP-6 is found in 80% of the *S. thermophilus* strains, with some harboring two occurrences of the defense systems, although manual inspection revealed that in many cases (183/361), this defense system may have been inactivated as it is present as pseudogenes (**Supplementary Data S5**).

Most of these systems have been validated as active against *S. thermophilus* phages (**Figure 5B**). In addition to the characterized CRISPR-Cas systems, we recently assessed the efficacy of many accessory defense systems, confirming activity for 17 out of 18 tested, with AbiJ being the only system that did not confer protection against the tested streptococcal phages^17^. Beyond RM systems, whose activity has not been systematically investigated in *S. thermophilus*, seven additional defense systems remain untested against streptococcal phages. These include four systems (VP1851, HEC-06, Retron type I-C, and TgvAB) highlighted in this study thanks to the latest DefenseFinder update and the expanded genome dataset; two systems (RloC and PrrC) whose activity depends on RM systems^54,55^, and Stk2^56^, which occurs exclusively as pseudogenes.

### *S. thermophilus* still shows potential as reservoir of novel defense systems

Using DefensePredictor, a recently developed machine-learning tool designed to evaluate the anti-phage defense potential of genes^7^, we further analyzed the same set of 364 *S. thermophilus* genomes to identify genes predicted to contribute to phage defense (**Figure 5C**). Across all genomes, 1.9% of the genes were predicted as defensive, with individual strains typically harboring between 1% and 2.7% of predicted defense genes in their genome (**Supplementary Figure S10C-D**). Of the 11,143 defensive genes identified, 78% matched validated defense systems, with the majority corresponding to CRISPR-Cas and RM systems (**Figure 5C**). These systems are both highly prevalent in *S. thermophilus* and typically encoded by multi-gene loci. Non-CRISPR and non-RM systems, including SthRAP systems, accounted for 9.1% of defense-associated genes and are typically present as accessory defense systems thus present in fewer strains (**Figure 5C**).

Among the 48 defense systems identified in *S. thermophilus*, seven systems (SthRAP-3, SthRAP-4, SthRAP-14, VP1851, SoFic, SEFIR, and Stk2) were never predicted as defensive by DefensePredictor in any of the strains harboring them (**Supplementary Data S6** and **Supplementary Figure S10E**). In addition, several systems were only partially detected by DefensePredictor (**Supplementary Figure S10E**). While partial detection often resulted from missing genes in multi-gene systems, in several cases complete systems were overlooked, such as SthRAP-6 (detected in only 5.1% of cases), AbiE (26.8%), and PrrC (20%). Interestingly, 21.9% of the identified genes did not correspond to validated defense systems and therefore represent additional candidate defense systems (**Figure 5C**). Furthermore, in most strains (78/81), the 32 GIs associated with phage defense were significantly enriched in these DefensePredictor candidate systems compared to the rest of the genome (Fisher’s exact test, Benjamini–Hochberg adjusted P < 0.05), further supporting their potential for uncovering novel defense systems in *S. thermophilus* (**Supplementary Data S7**).

## 2. Discussion

The analysis of the genomic localisation of defense systems has enabled the discovery of previously unknown defense mechanisms^3^. Although the examination of the gene neighborhood of known defense systems has been the main localisation-based strategy for uncovering new systems, recent studies have shown that defense systems are often located within specific genomic regions^23–25^. These regions consist of accessory genes flanked by core genes, which delineate loci whose content frequently varies from one strain to another. The presence of anti-phage defense systems within some of these regions motivated their designation as chromosomal defense hotspots and opened the way for the systematic investigation as promising reservoirs of uncharacterized defense systems.

In this study, we focused on *S. thermophilus*, a LAB widely used in the dairy industry. By analysing the genomic localisation of known defense systems in 81 complete genomes, we identified 30 GIs in which defense systems are commonly found, fewer than the 41 defense hotspots reported in *E. coli*^23^. However, the genome of *S. thermophilus* has undergone substantial gene loss associated with species domestication^30^, resulting in a smaller genome (ca. 1.8 Mb compared to 4.6 Mb in *E. coli*). Consequently, the proportion of GIs involved in phage defense is comparatively higher in *S. thermophilus*. Furthermore, plasmids are rare in *S. thermophilus* and, when present, are typically small (< 10 kb). As a result, defense systems are encoded almost exclusively on the chromosome^17^, which may contribute to the relatively high number of defense-associated GIs compared with plasmid-rich species. Our analysis also suggests that additional GIs may exist, as illustrated by two putative GIs (GI-A and GI-B) that contained only candidate defense systems, which are likely to be more clearly identified as more genomes are analyzed and new defense systems are uncovered. The relatively high number of GIs associated with phage defense in *S. thermophilus* may reflect the strong selective pressure imposed by virulent phages in the dairy environment, where only strains carrying adequate defense systems can persist. In addition, with few exceptions such as SthRAP-6 and Stk2, very few defense systems were found as pseudogenes (**Supplementary Data S5**), further supporting their importance for *S. thermophilus* survival in this environment.

The GIs exhibited substantial variation in their defense system content, with some being specific to particular defense systems (e.g., GI1 and GI5), while others such as GI11, GI12, and GI29 harbored a diverse anti-phage repertoire, which is expected to be even more diverse as more genomes are added. Their occupancy rates also varied greatly. In contrast to *E. coli*, where most GIs are occupied in only a few strains, *S. thermophilus* GIs were frequently occupied across many strains, and several contained defense systems in more than 75% of the genomes analysed, suggesting that they function as dedicated phage defense loci. In many strains, the GIs were occupied by genes unrelated to known defense systems. *In silico* prediction combined with manual inspection of GI content revealed that biosynthetic gene clusters and genes involved in metal resistance, for example, were also present in some GIs. These additional accessory functions motivated the consistent use of the term “genomic island” instead of chromosomal defense hotspots throughout the manuscript to reflect that the regions identified here are not necessarily entirely dedicated to phage defense.

We delineated GIs using a targeted strategy focused on defense systems, combining their chromosomal locations with a core-genome analysis to identify regions involved in phage defense. Unlike previous studies, we applied stringent criteria to define core genes, considering only orthogroups present in 100% of the genomes and relying exclusively on single-copy core genes to serve as the flanking boundaries of each GI. This approach makes it straightforward to locate a given GI across genomes based on a single conserved gene. However, it can also overestimate the number of GIs that appear to be occupied by genes unrelated to defense. This occurs when multi-copy core genes (such as transposases) or genes that have been inactivated in a few genomes cause their orthogroups to fall below the ubiquity threshold, leading to the selection of incorrect flanking genes.

Similarly to previous studies showing that defense systems often colocalize with MGEs ^22^, *S. thermophilus* GIs contained several ICEs, phage satellites, as well as mobility markers such as transposases and integrases. Prophages, on the other hand, are known to be rare in *S. thermophilus* and were not found integrated within the GIs, as defined in this study. In many cases, however, it was not possible to identify putative mobilization mechanisms. Although non-canonical forms of mobilization may be involved, *S. thermophilus* is also a naturally competent bacterium, enable to take up and integrate environmental DNA through transformation^57^. Furthermore, deletion of mobilome-associated genes may have occurred during evolution, leading to the fixation of GIs that provide advantageous adaptive functions^27^. Such deletions have been documented in *S. thermophilus* ICEs, where the loss of conjugation, recombination, and regulation modules has given rise to certain Integrative and Mobilizable Elements (IMEs)^58^.

Leveraging these GIs, we aimed at identifying novel defense systems. By focusing on three GIs, we uncovered 15 defense systems that were previously unknown in *S. thermophilus*, which we named SthRAP-1-15. While five systems appeared to be homologs or putative new variants of previously described defense systems, ten SthRAP systems represent entirely new defense systems with potentially unique modes of action. Prevalence analysis showed they were not restricted to *S. thermophilus* but were instead broadly distributed across the *Bacillota* phylum and, in most cases, even more distantly related phyla, although they typically occurred in fewer than 1% of the genomes analyzed, consistent with previous observations for defense systems^53^. Of note, two systems, SthRAP-10 and SthRAP-13 were only found in the *Streptococcus* genus suggesting that some systems might be specific to some bacterial genera, further supporting investigation of anti-defense mechanisms in non-model bacteria. Machine-learning tools recently developed to predict defense systems^7,8^ are likely to become the most powerful strategy for identifying novel defense mechanisms. However, applying DefensePredictor to our *S. thermophilus* dataset revealed that several defense systems, including three newly discovered ones (SthRAP-3, SthRAP-4, and SthRAP-14), were not predicted as defensive in any of the genomes examined. Other systems were only partially predicted, with only one gene identified as defensive, or predicted in some genomes but not others. These observations underscore those complementary strategies, such as investigating GIs enriched in defense systems, are still essential for fully characterizing the antiviral arsenal of certain bacteria.

The identification of the SthRAP systems expands the repertoire of defense mechanisms that can be leveraged to strengthen phage resistance in industrial *S. thermophilus* strains^17^. This includes rare systems such as SthRAP-1, displaying highly efficiency and which may be less susceptible to phage counter-adaptation due to their limited occurrence and, consequently, reduced exposure to phages. In addition to the transfer of individual defense systems between strains to modulate phage resistance, the GIs identified here also show strong potential for industrial applications. Entire GIs could potentially be exchanged between strains by amplifying the full regions, including the conserved flanking genes, which would allow their transfer through natural competence and thus enable the generation of non-GMO strains. Conversely, empty GIs present in industrial strains could serve as dedicated integration sites for defense systems, as these loci naturally harbor such systems in other *S. thermophilus* genomes. Beyond their industrial relevance, *S. thermophilus* GIs also represent promising reservoirs for the discovery of additional, previously uncharacterized defense systems.

## 3. Methods

### Genomic island delineation

Complete *S. thermophilus* genomes from the NCBI Reference Sequence (RefSeq) database were downloaded in March 2023. Out of the 86 genomes, five (GCF_028463905.1, GCF_903886595.1, GCF_015190465.1, GCF_903886745.1, GCF_022024355.1) were excluded due to assembly issues (**Supplementary Data S8**). To ensure consistency, all genomes were re-annotated using Bakta^59^ with default parameters.

Defense systems were detected using DefenseFinder v1.2.2 (models v1.2.3)^53^ and PADLOC v2.0.0 (models v2.0.0)^28^. Incomplete defense systems labelled as “other”, “cas adaptation”, or “cas_cluster” were excluded. Predictions from both tools were merged using a custom script. Following manual curation, Viperin_solo and PD-T4-6 were excluded from the analysis due to mispredictions. Closely related systems with overlapping gene predictions (i.e., PD-T7-2 and Gao19, as well as Abi2 and AbiD) were combined into single systems (i.e., Gao19 and AbiD, respectively). When one gene was identified as part of both a candidate (identified by PADLOC) and a validated defense system (i.e., PDC-S04 and PDC-S06 matched RloC and SoFic, respectively), only the annotation corresponding to the validated defense systems was kept. A full list of the identified defense systems is provided in **Supplementary Data S9**. Additionally, DefensePredictor^7^ was used to identify putative defensive genes using a log-odds cutoff of 7.2. Enrichment of candidate defense genes (i.e., genes predicted to be defensive but excluding those belonging to experimentally validated systems identified by DefenseFinder or PADLOC, as well as SthRAP systems) within GIs was assessed using Fisher’s exact tests in R, with Benjamini–Hochberg correction for multiple testing (adjusted P < 0.05).

For the core genome analysis, plasmids were filtered out from the dataset to focus on bacterial chromosomes. Core genes, defined as those present in 100% of the genomes, were identified using OrthoFinder^60^. Additionally, the type strain of *Streptococcus vestibularis* (GCF_900636445.1) was used as an outgroup for the single-copy core protein phylogeny inferred with OrthoFinder.

To define GIs associated with phage defense, the genomic context of defense systems identified in each strain was analyzed using a custom R script to identify the nearest upstream and downstream single-copy core genes, which were designated as GI boundaries (**Figure 1A**). The resulting GIs were then mapped onto the 81 genomes including that of strain DGCC7710 which was usedas a reference *S. thermophilus* genome for visualization. Information about flanking genes for each GI in the reference genome can be found in **Supplementary Data S1**.

### MGE prediction and biosynthetic gene clusters

MGEs associated with GIs were identified using a combination of tools and manual curation. Genomes were annotated with mobileOG-db v1.0 to detect genes related to the mobilome^61^. geNomad^62^ was then used to identify phage genes within genomic islands, which were compared with the manual curation of phage satellites (PS) in *S. thermophilus*^32^. This confirmed that the detected phage genes corresponded to PSs rather than prophages, the latter being rare in *S. thermophilus*. ICEs were identified manually by searching for genes involved in conjugative transfer. In GI lacking PSs or ICEs, the presence of transposase or integrase genes was assessed using Bakta annotations. Genomes were also searched for biosynthetic gene clusters using antiSMASH 8.0^63^.

### Bacteria growth conditions

Strains, phages, plasmids, and primers used in this study are listed in **Supplementary Data S10** and **Supplementary Data S11**. *E. coli* was grown overnight (O/N) at 37 °C with shaking in Brain Heart Infusion (BHI; Difco) medium. *S. thermophilus* was cultured O/N without shaking in 0.5x M17 medium (Nutri-Bact) supplemented with 0.5% (w/v) lactose (LM17) at 37 °C. For transformation experiments, media were solidified with 1.5% agar (Laboratoire Mat) and supplemented with erythromycin at final concentrations of 150 µg/ml for *E. coli* and 2.5 µg/ml for *S. thermophilus*. Plates were incubated at 37 °C for *E. coli* and 42 °C for *S. thermophilus*.

### Candidate defense system cloning

Candidate defense systems (or individual defense genes) were obtained either by PCR from bacterial strains in our collection using the Q5 High-Fidelity DNA Polymerase (New England Biolabs) or synthesized by Bio Basic, GenScript, or Integrated DNA Technologies. PCR products were cloned into the low-copy vector pTRKL2^64^ using Gibson assembly, and the resulting reaction mixtures were used to transform chemically competent *E. coli* NEB5α cells. Positive clones were identified by PCR screening with Taq DNA polymerase (Bio Basic) and confirmed by Sanger sequencing. Plasmid constructs were extracted from *E. coli* using the QIAprep Spin Miniprep Kit (Qiagen) and introduced into *S. thermophilus* strains lacking the corresponding candidate systems via either electroporation or natural competence, as described previously^17^. The presence and correct sequence of the candidate systems in the transformants were verified by PCR screening and Sanger sequencing.

### Assessment of defense system efficiency

The activity of the candidate defense systems was assessed using spot test assays with phage lysates at concentrations of approximately 10⁸–10⁹ PFU/ml (three biological replicates). Modified overlay (0.4% agarose, 0.5% glycine, and 0.1% milk) supplemented with 10 mM CaCl₂was mixed with 15% of an O/N bacterial culture and poured onto LM17 plates (1% agar) containing erythromycin and 10 mM CaCl₂. Next, 3 μl drops of tenfold serial dilutions of each phage lysate were spotted onto the surface and allowed to dry before incubation. Phage titers were calculated following O/N incubation of the plates. The efficiency of plaquing (EOP) was calculated as the ratio of the phage titer in the presence of the defense system to that obtained with the strain carrying the empty vector.

The activity of validated defense systems was further assessed using liquid assays. O/N cultures of *S. thermophilus* strains carrying either the empty pTRKL2 vector or a defense system were diluted to an OD₆₀₀ of 0.2 in filtered LM17 supplemented with 10 mM CaCl₂. A volume of 20 µl of this suspension was added to wells of a 96-well plate containing 160 µl of LM17 with 10 mM CaCl₂, followed by the addition of 20 µl of phage buffer or tenfold serially diluted phage lysate to test different multiplicities of infection (MOIs). Plates were incubated at 37 °C for 16 hours in a plate reader and OD₆₀₀ measurements were taken every 10 minutes after 10 seconds of orbital shaking. Each growth curve represents the average of three biological replicates.

### Functional and structural analysis

Sequence-based functional annotation was performed using InterProScan^65^ to identify conserved domains, DeepTMHMM^66^ to predict transmembrane topology, and HHpred (https://toolkit.tuebingen.mpg.de/tools/hhpred) to detect remote homology. Structural predictions were generated with AlphaFold3 (AF3)^67^. Predicted multimerization was assessed, with ipTM scores above 0.6 interpreted as supporting multimer formation. For two-component defense systems, we also evaluated the likelihood of heterodimer formation (ipTM > 0.6). Predicted interactions with nucleic acids or ligands (when relevant) were also examined, and ipTM scores above 0.8 were considered as indicative of a probable interaction. Detailed ipTM and pTM values are provided in **Supplementary Data S3**. AF3 monomeric models were submitted to the Dali server^68^ to identify potential structural homologs. Structural visualizations were prepared using PyMOL and structure superimpositions were performed with cealign.

### Defense system prevalence assessment

The prevalence of defense systems in bacterial genomes was assessed using MacSyFinder v2^69^. HMMs were obtained from existing databases (PFAM or PANTHER) for defense proteins with conserved domains spanning the full protein length or from PADLOC for defense systems (SthRAP-5, -11, -14) corresponding to candidate systems (PDC-S50, PDC-M01, and PDC-S34). Custom HMMs were generated for proteins lacking existing profiles. To do so, homologous proteins were retrieved using BLASTp (≥50% coverage; e-value ≤ 1e-5) against a RefSeq database of complete bacterial genomes (June 2025). Identical sequences were removed with SeqKit2^70^. For protein families with more than 1,000 sequences, MMseqs2^71^ was used to cluster sequences at 30% identity and 80% coverage, retaining only clusters containing at least five sequences. Redundancy in clusters with over 100 sequences was further reduced using CD-HIT (90% identity; 90% coverage). Multiple sequence alignments were generated using MUSCLE v5.1.0^72^ (>200 sequences) or MAFFT v7.526^73^ (<200 sequences). HMMs were then built for each cluster using HMMER v3.4^74^ with default settings.

For each defense system, model definitions were established following MacSyFinder guidelines, allowing up to five gene insertions for two-gene systems, similar to the rules implemented in DefenseFinder. Defense systems were first searched with MacSyFinder in the complete *S. thermophilus* genomes to validate model quality. A GA score threshold of 18 was set to the PADLOC HMM for SthRAP-11B to enable the detection of all occurrences. The defense systems were then searched across all *S. thermophilus* genomes (N = 364, June 2025) and complete bacterial genomes (N = 50,676, RefSeq June 2025). For single-gene defense systems, score distributions were examined, and low-score hits were manually inspected by verifying conserved domains and/or by predicting protein structures with AF3 to remove false positives. For each homolog identified, the taxonomic information of the bacterium was retrieved using its TaxID and the NCBITaxa module from the ETE3 Python package^75^. Defense prevalence within each bacterial class or family was calculated as the log_10_ of the number of occurrences in a given taxon divided by the total number of genomes belonging to that taxon. The latest version of DefenseFinder (v2.0.0, models v2.0.2)^53^ was also used to predict known defense systems in *S. thermophilus* genomes for **Figure 5B**. For the prevalence of VP1848 and Septu type I, the HMMs and system definitions were retrieved from DefenseFinder.

## 5. Funding

AL was supported by a postdoctoral fellowship from Wallonie-Bruxelles International under the grant SUB/2022/554282. VS was supported by SNSF Postdoc.Mobility P500PB_214419 and SNF Return Grant P5R5PB_230730. SM acknowledges funding from NSERC (Discovery Program). SM is a member of the strategic networks PROTEO and Op+Lait of the FRQNT and previously held the Canada Research Chair in Bacteriophages (2011–2025).

## 6. Competing interest

Patent applications were filled in the context of this work.

## References

1. Hampton, H. G., Watson, B. N. J. & Fineran, P. C. The arms race between bacteria and their phage foes. Nature 577, 327–336 (2020).

2. Leprince, A. & Mahillon, J. Phage adsorption to Gram-positive bacteria. Viruses 15, 196 (2023).

3. Georjon, H. & Bernheim, A. The highly diverse antiphage defence systems of bacteria. Nat. Rev. Microbiol. 21, 686–700 (2023).

4. Hu, H. et al. Structure and mechanism of the Zorya anti-phage defense system. Nature 639, 1093–1101 (2024).

5. Tuck, O. T. et al. Genome integrity sensing by the broad-spectrum Hachiman antiphage defense complex. Cell 187, 6914–6928 (2024).

6. Zhang, Z. et al. Kiwa is a membrane-embedded defense supercomplex activated at phage attachment sites. Cell 188, 5862–5877 (2025).

7. DeWeirdt, P. C., Mahoney, E. M. & Laub, M. T. DefensePredictor: A machine learning model to discover prokaryotic immune systems. Science 392, eadv7924 (2026).

8. Mordret, E. et al. Protein and genomic language models uncover the unexplored diversity of bacterial immunity. Science 392, eadv8275 (2026).

9. Samson, J. E. & Moineau, S. Bacteriophages in food fermentations: new frontiers in a continuous arms race. Annu. Rev. Food Sci. Technol. 4, 347–368 (2013).

10. Jolicoeur, A. P. et al. Longitudinal study of *Lactococcus* phages in a Canadian cheese factory. Appl. Environ. Microbiol. 89, e0042123 (2023).

11. Romero, D. A., Magill, D., Millen, A., Horvath, P. & Fremaux, C. Dairy lactococcal and streptococcal phage-host interactions: an industrial perspective in an evolving phage landscape. FEMS Microbiol. Rev. 44, 909–932 (2020).

12. Philippe, C. et al. The never-ending battle between lactic acid bacteria and their phages. FEMS Microbiol. Rev. 47, fuad035 (2023).

13. Chopin, M.-C., Chopin, A. & Bidnenko, E. Phage abortive infection in lactococci: variations on a theme. Curr. Opin. Microbiol. 8, 473–479 (2005).

14. Leprince, A. & Moineau, S. CRISPR-Cas in the cheese industry. Annu. Rev. Virol. 12, 491–512 (2025).

15. Grafakou, A. et al. Discovery of antiphage systems in the lactococcal plasmidome. Nucleic Acids Res. 52, 9760–9776 (2024).

16. Kelleher, P. et al. Phage defence loci of *Streptococcus thermophilus*-tip of the anti-phage iceberg? Nucleic Acids Res. 52, 11853–11869 (2024).

17. Leprince, A. et al. Strengthening phage resistance of *Streptococcus thermophilus* by leveraging complementary defense systems. Nat. Commun. 16, 7142 (2025).

18. Makarova, K. S., Wolf, Y. I., Snir, S. & Koonin, E. V. Defense islands in bacterial and archaeal genomes and prediction of novel defense systems. J. Bacteriol. 193, 6039–6056 (2011).

19. Doron, S. et al. Systematic discovery of antiphage defense systems in the microbial pangenome. Science 359, (2018).

20. Gao, L. et al. Diverse enzymatic activities mediate antiviral immunity in prokaryotes. Science 369, 1077–1084 (2020).

21. Rousset, F. et al. Phages and their satellites encode hotspots of antiviral systems. Cell Host Microbe 30, 740–753 (2022).

22. Beamud, B., Benz, F. & Bikard, D. Going viral: The role of mobile genetic elements in bacterial immunity. Cell Host Microbe 32, 804–819 (2024).

23. Hochhauser, D., Millman, A. & Sorek, R. The defense island repertoire of the *Escherichia coli* pan-genome. PLoS Genet. 19, e1010694 (2023).

24. Johnson, M. C. et al. Core defense hotspots within *Pseudomonas aeruginosa* are a consistent and rich source of anti-phage defense systems. Nucleic Acids Res. 51, 4995–5005 (2023).

25. Yi, W. et al. In silico characterization of defense system hotspots in *Acinetobacter* spp. *Commun*. Biol. 8, 39 (2025).

26. Rodriguez-Valera, F., Martin-Cuadrado, A.-B. & López-Pérez, M. Flexible genomic islands as drivers of genome evolution. Curr. Opin. Microbiol. 31, 154–160 (2016).

27. Bellanger, X., Payot, S., Leblond-Bourget, N. & Guédon, G. Conjugative and mobilizable genomic islands in bacteria: evolution and diversity. FEMS Microbiol. Rev. 38, 720–760 (2014).

28. Payne, L. J. et al. Identification and classification of antiviral defence systems in bacteria and archaea with PADLOC reveals new system types. Nucleic Acids Res. 49, 10868–10878 (2021).

29. Payne, L. J., Hughes, T. C. D., Fineran, P. C. & Jackson, S. A. New antiviral defences are genetically embedded within prokaryotic immune systems. bioRxiv (2024) doi:10.1101/2024.01.29.577857.

30. Somerville, V. et al. Genomic and phenotypic imprints of microbial domestication on cheese starter cultures. Nat. Commun. 15, 8642 (2024).

31. Burrus, V., Bontemps, C., Decaris, B. & Guédon, G. Characterization of a novel type II restriction-modification system, Sth368I, encoded by the integrative element ICESt1 of *Streptococcus thermophilus* CNRZ368. Appl. Environ. Microbiol. 67, 1522–1528 (2001).

32. Morency, C., Rousseau, G. M., Morneau, Z. & Moineau, S. Phage satellites induced by virulent phages are mobilized by natural competence leading to phage resistance in a new host. Nat. Commun. 1–18 (2026).

33. Philippe, C. et al. Novel genus of phages infecting Streptococcus thermophilus: Genomic and morphological characterization. Appl. Environ. Microbiol. 86, e00227–20 (2020).

34. Aframian, N. & Eldar, A. Abortive infection antiphage defense systems: separating mechanism and phenotype. Trends Microbiol. 31, 1003–1012 (2023).

35. Andersen, S. E. et al. Small serine recombinases are markers for antiphage defense system discovery. PLoS Biol. 24, e3003991 (2026).

36. Li, M. et al. Codon-usage-based inhibition of HIV protein synthesis by human schlafen 11. Nature 491, 125–128 (2012).

37. Yang, J.-Y. et al. Structure of Schlafen13 reveals a new class of tRNA/rRNA-targeting RNase engaged in translational control. Nat. Commun. 9, 1165 (2018).

38. Metzner, F. J. et al. Mechanistic understanding of human SLFN11. Nat. Commun. 13, 5464 (2022).

39. Perez Taboada, V., et al. Bacterial Schlafen proteins mediate phage defence. Nat. Microbiol. 11, 1037–1048 (2026).

40. Aravind, L., Iyer, L. M., Leipe, D. D. & Koonin, E. V. A novel family of P-loop NTPases with an unusual phyletic distribution and transmembrane segments inserted within the NTPase domain. Genome Biol. 5, R30 (2004).

41. Li, Y. et al. PtuA and PtuB assemble into an inflammasome-like oligomer for anti-phage defense. Nat. Struct. Mol. Biol. 31, 413–423 (2024).

42. Sabonis, D., et al. TIR domains produce histidine-ADPR conjugates as immune signaling molecules in bacteria. bioRxiv (2024) doi:10.1101/2024.01.03.573942.

43. Shi, Y. et al. Structural characterization of macro domain-containing Thoeris antiphage defense systems. Sci. Adv. 10, eadn3310 (2024).

44. Groves, M. R., Hanlon, N., Turowski, P., Hemmings, B. A. & Barford, D. The structure of the protein phosphatase 2A PR65/A subunit reveals the conformation of its 15 tandemly repeated HEAT motifs. Cell 96, 99–110 (1999).

45. Kschonsak, M. et al. Structural basis for a safety-belt mechanism that anchors condensin to chromosomes. Cell 171, 588–600 (2017).

46. Getz, L. J., Fairburn, S. R., Vivian Liu, Y., Qian, A. L. & Maxwell, K. L. Integrons are anti-phage defence libraries in *Vibrio parahaemolyticus*. Nat. Microbiol. 10, 724–733 (2025).

47. Tal, N. et al. Bacteria deplete deoxynucleotides to defend against bacteriophage infection. Nat. Microbiol. 7, 1200–1209 (2022).

48. Ibryashkina, E. M., Sasnauskas, G., Solonin, A. S., Zakharova, M. V. & Siksnys, V. Oligomeric structure diversity within the GIY-YIG nuclease family. J. Mol. Biol. 387, 10–16 (2009).

49. Kuznetsova, E. et al. Functional diversity of haloacid dehalogenase superfamily phosphatases from *Saccharomyces cerevisiae*: biochemical, structural, and evolutionary insights. J. Biol. Chem. 290, 18678–18698 (2015).

50. Hadži, S. et al. Ribosome-dependent *Vibrio cholerae* mRNAse HigB2 is regulated by a β-strand sliding mechanism. Nucleic Acids Res. 45, 4972–4983 (2017).

51. Zhou, Q. et al. Architecture of the synaptotagmin-SNARE machinery for neuronal exocytosis. Nature 525, 62–67 (2015).

52. Posner, M. G. et al. Distinctive phosphoinositide- and Ca2+-binding properties of normal and cognitive performance-linked variant forms of KIBRA C2 domain. J. Biol. Chem. 293, 9335–9344 (2018).

53. Tesson, F. et al. Systematic and quantitative view of the antiviral arsenal of prokaryotes. Nat. Commun. 13, 2561 (2021).

54. Davidov, E. & Kaufmann, G. RloC: a wobble nucleotide-excising and zinc-responsive bacterial tRNase. Mol. Microbiol. 69, 1560–1574 (2008).

55. Penner, M., Morad, I., Snyder, L. & Kaufmann, G. Phage T4-coded Stp: double-edged effector of coupled DNA and tRNA-restriction systems. J. Mol. Biol. 249, 857–868 (1995).

56. Depardieu, F. et al. A eukaryotic-like serine/threonine kinase protects staphylococci against phages. Cell host & microbe 20, 471–481 (2016).

57. Di Giacomo, S. et al. Expanding natural transformation to improve beneficial lactic acid bacteria. FEMS Microbiol. Rev. 46, fuac014 (2022).

58. Pavlovic, G., Burrus, V., Gintz, B., Decaris, B. & Guédon, G. Evolution of genomic islands by deletion and tandem accretion by site-specific recombination: ICESt1-related elements from *Streptococcus thermophilus*. Microbiology 150, 759–774 (2004).

59. Schwengers, O. et al. Bakta: rapid and standardized annotation of bacterial genomes via alignment-free sequence identification. Microb. Genom. 7, 000685 (2021).

60. Emms, D. M. & Kelly, S. OrthoFinder: phylogenetic orthology inference for comparative genomics. Genome Biol. 20, 238 (2019).

61. Brown, C. L. et al. MobileOG-db: A manually curated database of protein families mediating the life cycle of bacterial mobile genetic elements. Appl. Environ. Microbiol. 88, e0099122 (2022).

62. Camargo, A. P. et al. Identification of mobile genetic elements with geNomad. Nat. Biotechnol. 42, 1303–1312 (2024).

63. Blin, K. et al. antiSMASH 8.0: extended gene cluster detection capabilities and analyses of chemistry, enzymology, and regulation. Nucleic Acids Res. 53, W32–W38 (2025).

64. O’Sullivan, D. J. & Klaenhammer, T. R. High- and low-copy-number *Lactococcus* shuttle cloning vectors with features for clone screening. Gene 137, 227–231 (1993).

65. Blum, M. et al. InterPro: the protein sequence classification resource in 2025. Nucleic Acids Res. 53, D444–D456 (2025).

66. Hallgren, J., et al. DeepTMHMM predicts alpha and beta transmembrane proteins using deep neural networks. bioRxiv (2022) doi:10.1101/2022.04.08.487609.

67. Abramson, J. et al. Accurate structure prediction of biomolecular interactions with AlphaFold 3. Nature 630, 493–500 (2024).

68. Holm, L., Laiho, A., Törönen, P. & Salgado, M. DALI shines a light on remote homologs: One hundred discoveries. Protein Sci. 32, e4519 (2023).

69. Néron, B. et al. MacSyFinder v2: Improved modelling and search engine to identify molecular systems in genomes. Peer Community J. 3, (2023).

70. Shen, W., Sipos, B. & Zhao, L. SeqKit2: A Swiss army knife for sequence and alignment processing. Imeta 3, e191 (2024).

71. Steinegger, M. & Söding, J. MMseqs2 enables sensitive protein sequence searching for the analysis of massive data sets. Nat. Biotechnol. 35, 1026–1028 (2017).

72. Edgar, R. C. Muscle5: High-accuracy alignment ensembles enable unbiased assessments of sequence homology and phylogeny. Nat. Commun. 13, 6968 (2022).

73. Katoh, K., Rozewicki, J. & Yamada, K. D. MAFFT online service: multiple sequence alignment, interactive sequence choice and visualization. Brief. Bioinform. 20, 1160–1166 (2019).

74. Eddy, S. R. Accelerated profile HMM searches. PLoS Comput. Biol. 7, e1002195 (2011).

75. Huerta-Cepas, J., Serra, F. & Bork, P. ETE 3: Reconstruction, analysis, and visualization of phylogenomic data. Mol. Biol. Evol. 33, 1635–1638 (2016).

