## Supplementary Figures for "Uncovering novel phage defense systems through genomic island analysis in *Streptococcus thermophilus*"

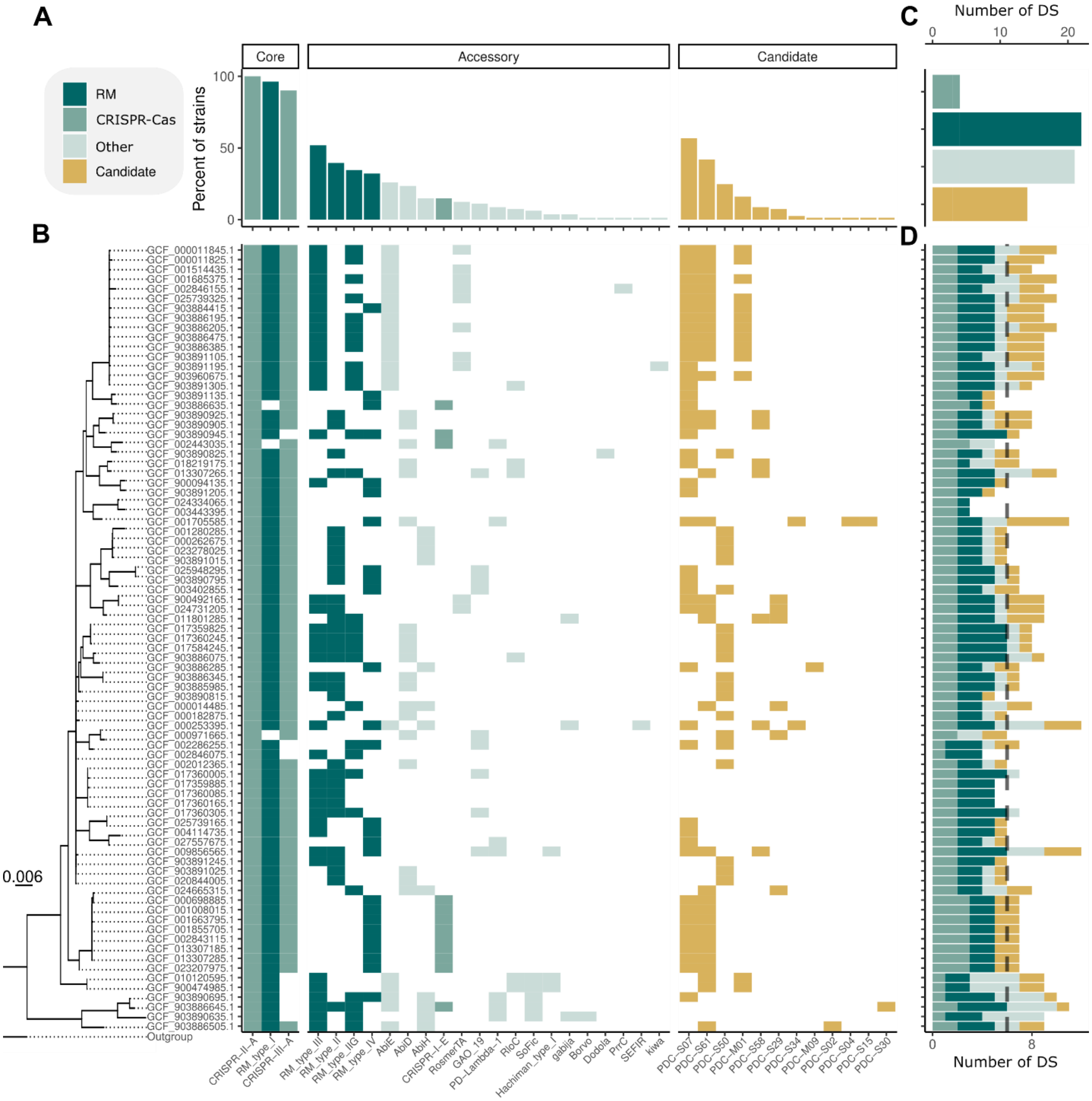


**Supplementary Figure S1 | Phylogenetic distribution and prevalence of defense systems in the 81 genomes of *S. thermophilus*. A.** Prevalence of defense systems predicted by DefenseFinder and PADLOC classified as core (>70% of genomes), accessory (<70%), or candidate (PADLOC predictions not yet experimentally validated). **B.** Distribution of defense systems per strain. The phylogenetic tree illustrates strain relationships, with branch lengths representing amino acid substitutions per site. **C.** Number of distinct defense systems or types per defense category. The category "Other" includes experimentally validated systems that are neither RM nor CRISPR-Cas. **D.** Number of defense systems per strain, with the dashed line indicating the dataset mean. DS, defense systems.

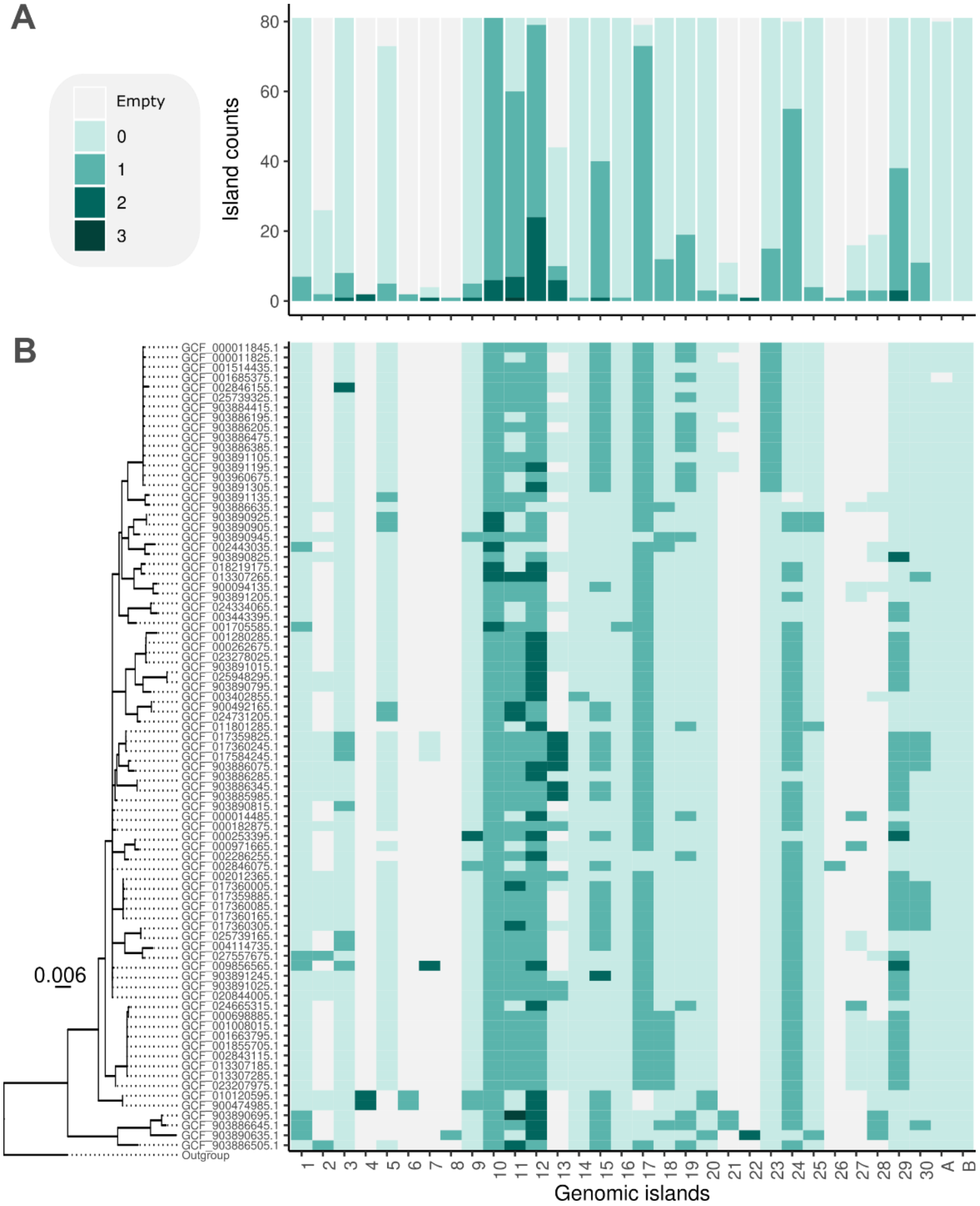


**Supplementary Figure S2 | Defense occupation of Genomic Islands (GIs) in the 81 *S. thermophilus* genomes.** Occupation status of the 30 validated GIs and the two candidate GIs based on defense system content. "Empty" indicates no genes between the two flanking core host genes. Blue shading denotes GIs occupied by genes: 0 means no known defense system is present, while 1, 2, or 3 indicate the number of experimentally validated defense systems within that GI (which excludes the candidate defense systems from PADLOC). **A.** Summary plot showing the count of each occupation category per GI. **B.** Occupation status of all GIs across strains, with the phylogenetic tree illustrating strain relationships (branch lengths represent amino acid substitutions per site).

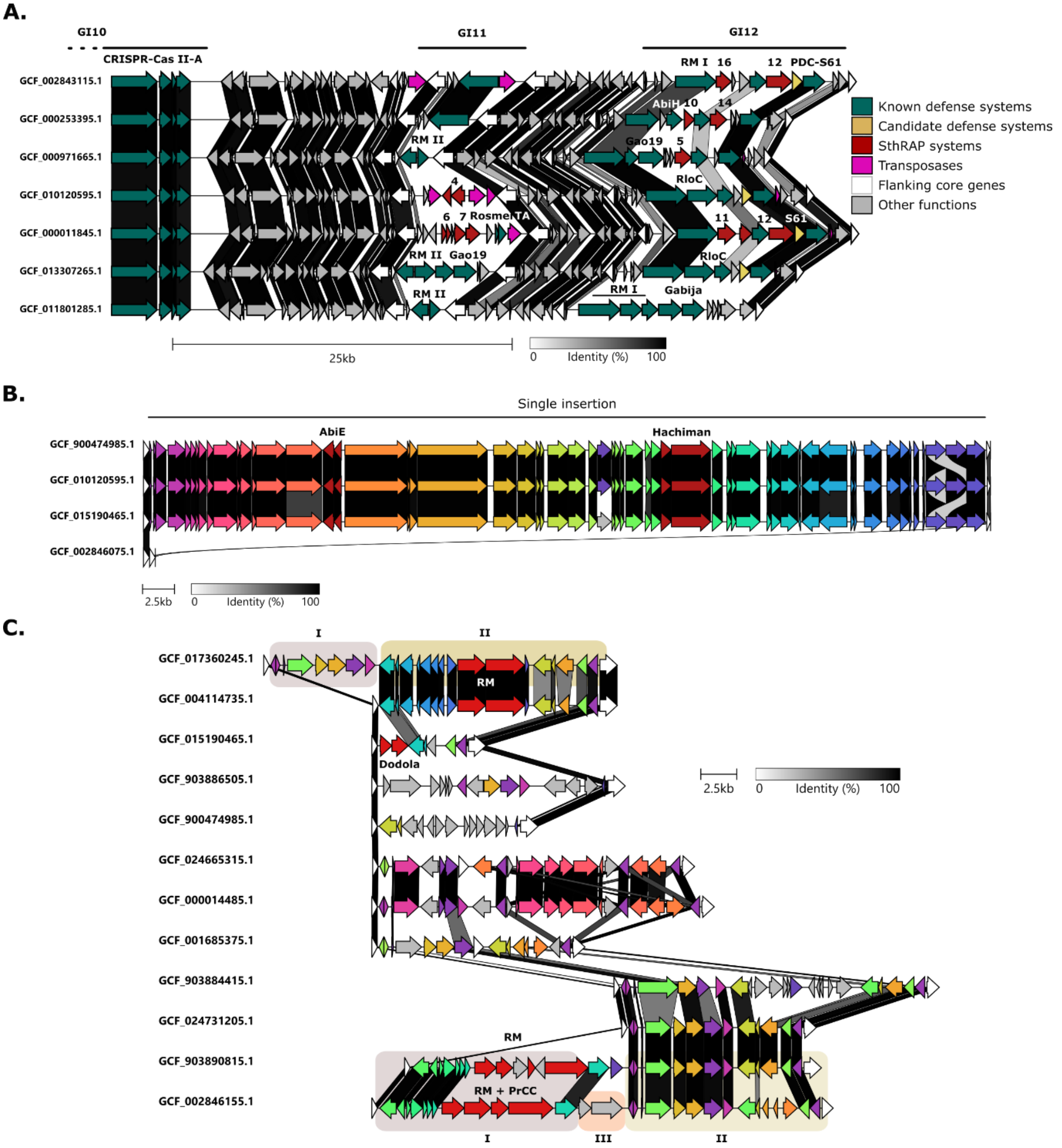


**Supplementary Figure S3 | Alignment of representative GIs associated with phage defense systems. A.** Alignment of representative genomes showing the region encompassing CRISPR–Cas type II-A (CRISPR1), GI11, and GI12 as well as illustrating examples of the diversity of defense systems found within these GIs. **B.** Alignment of GI4 for strains in which it is occupied. This shows an example of GI4 where a single large insertion is found. **C.** Representative alignment of GI29, which displays variable gene content between strains and illustrates multiple insertions within the same GI. Known defense systems are colored in red, while other genes are colored based on homology; genes sharing the same color encode deduced proteins with at least 30% amino acid identity. Distinct insertion events within the same GI are highlighted by squares labeled I, II, or III. Alignments were generated using clinker[1].

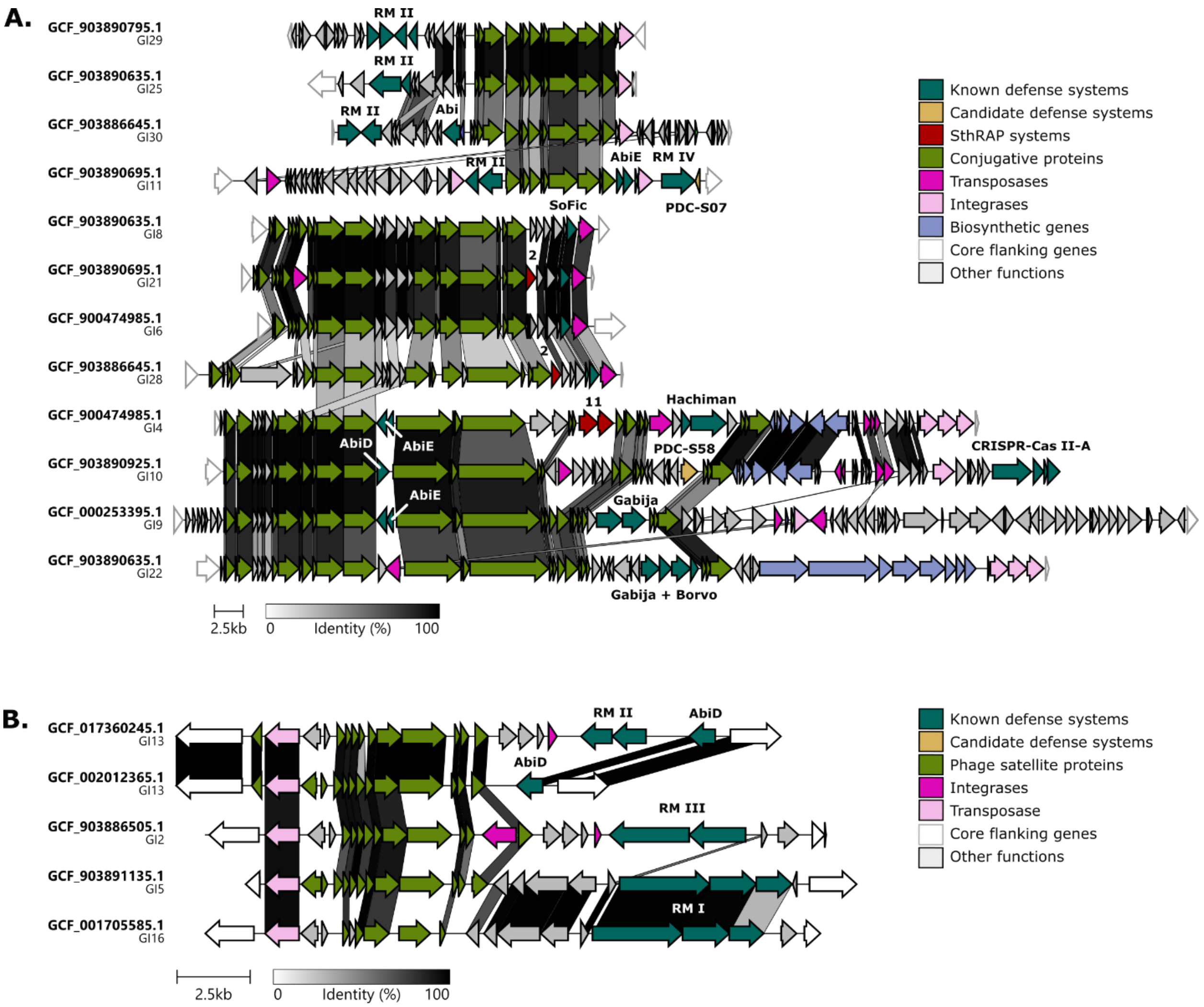


**Supplementary Figure S4 | Alignment of representative GIs harboring mobile genetic elements (MGEs) found in *S. thermophilus*.** The genomes and GIs from which the MGEs were extracted are indicated on the left. The links between the genes indicate amino acid identity of the encoded protein. The alignments were generated with clinker[1]. **A.** Integrative and Conjugative Elements (ICEs) and **B.** Phage satellites.

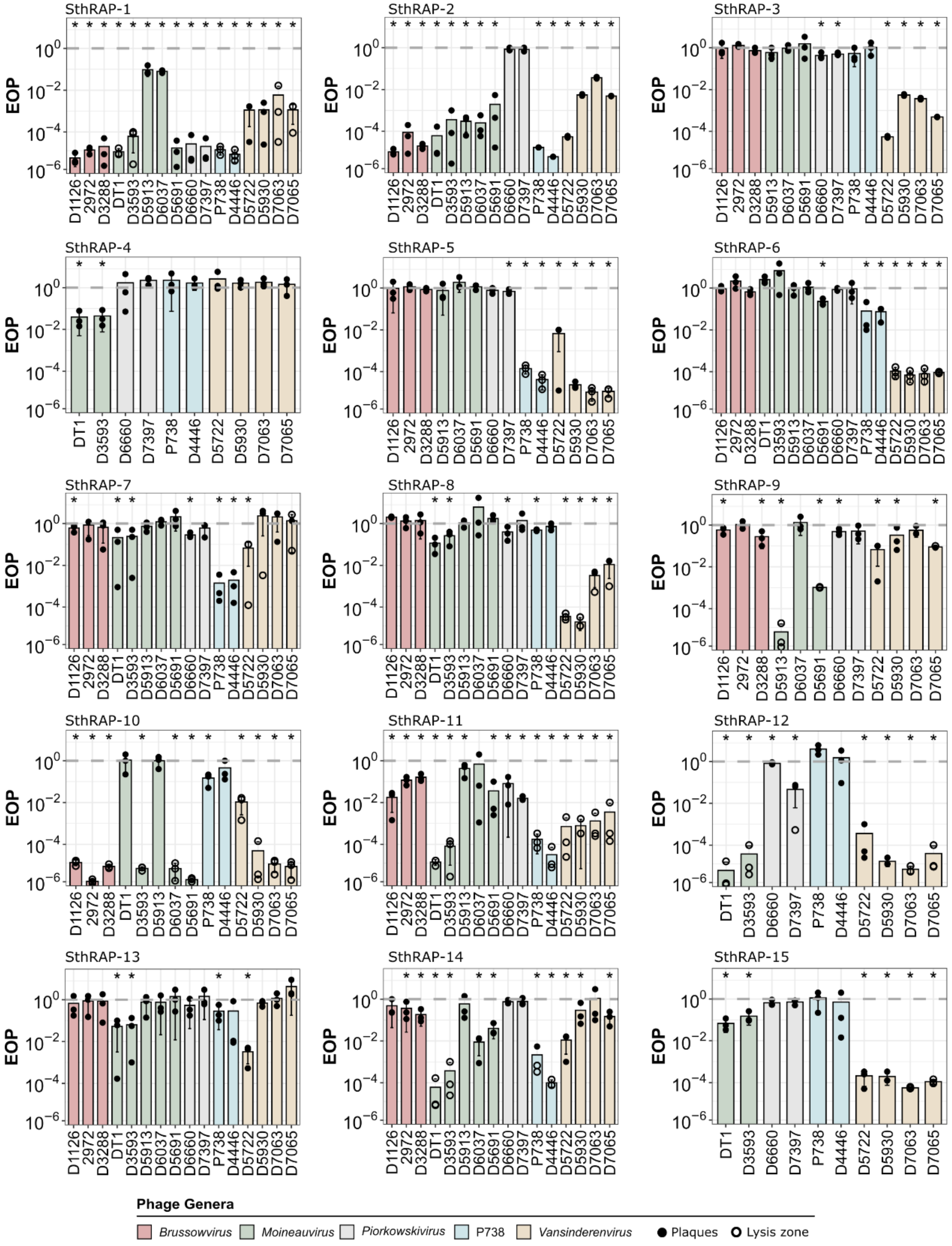


**Supplementary Figure S5 | Efficiency of plaquing (EOP) for each SthRAP system against *S. thermophilus* virulent phages.** Bars represent the mean EOP from three independent replicates, with individual values shown as circles. Filled circles indicate countable plaques, while hollow circles denote zones of lysis without distinct plaques. The dotted line at EOP = 1 corresponds to no anti-phage activity. Asterisks indicate statistically significant differences from EOP = 1 (t-test, $p < 0.05$). EOP, Efficiency of plaquing.

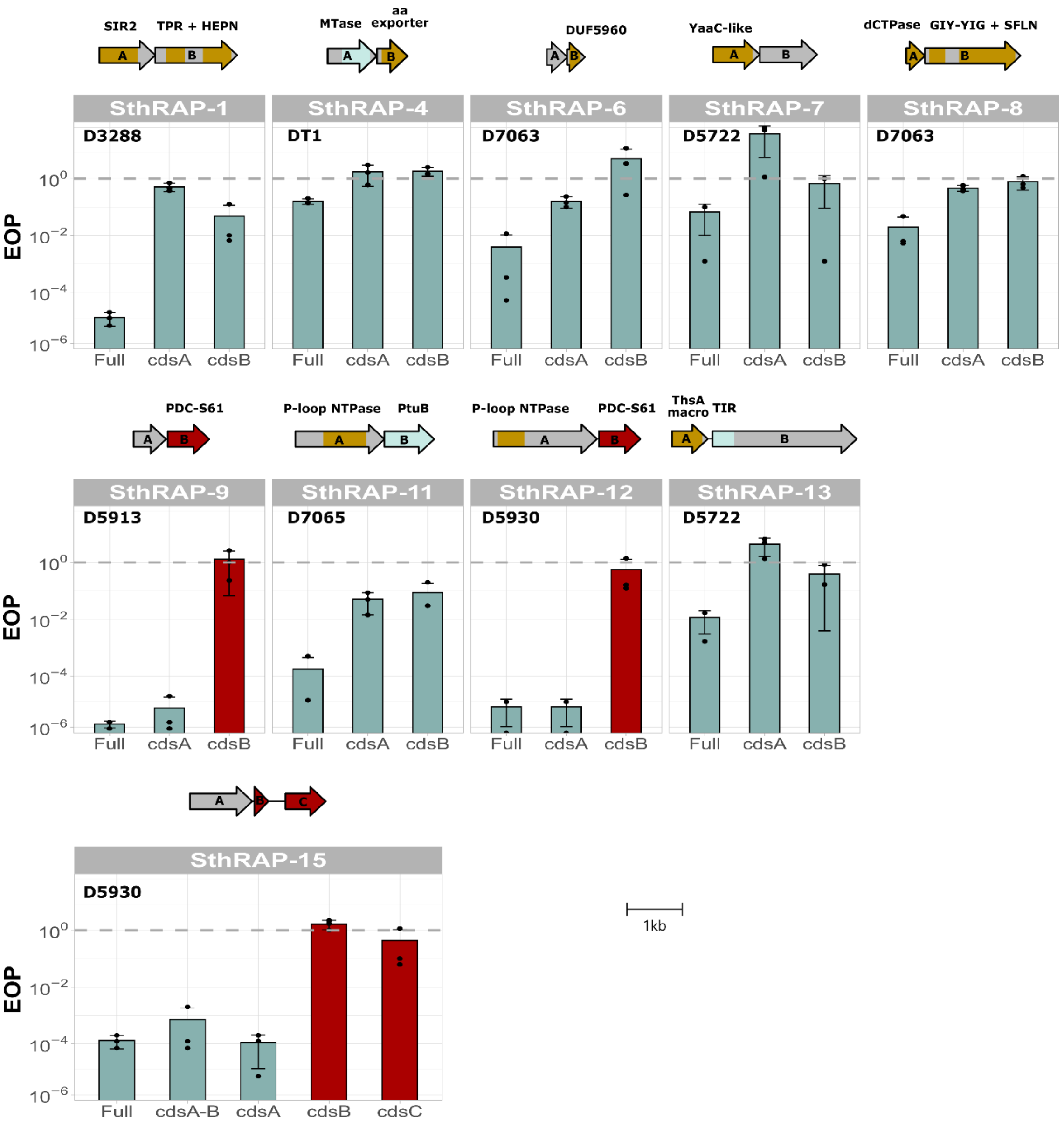


**Supplementary Figure S6 | Confirmation of multi-gene systems.** The contribution of individual genes to phage resistance was assessed by testing each gene separately. Genetic organization of each system is shown above the graphs, with conserved domains or structural folds highlighted. For SthRAP-9, SthRAP-12, and SthRAP-15, initial loci included genes that did not confer resistance and were excluded from the final system (highlighted in red). PDC-S61 indicates that the *cdsB* associated with SthRAP-9 and SthRAP-12 (as well as SthRAP-14 but not tested) was predicted as a standalone candidate by PADLOC. Tested phages are indicated in the upper part of each graph. Abbreviations: aa, Amino acid; dCTPase, dCTP pyrophosphatase; DUF, Domain of unknown function; GIY-YIG, GIY-YIG endonuclease; HEPN, Higher eukaryotes and prokaryotes nucleotide-binding; MTase, Methyltransferase; NTPase, Nucleoside-triphosphatase; PtuB, Septu subunit B; SFLN, Schlafen; SIR2, Sirtuin; ThsA, Thoeris subunit A; TIR, Toll/interleukin-1 receptor/resistance protein; TPR, Tetratricopeptide.

Internal

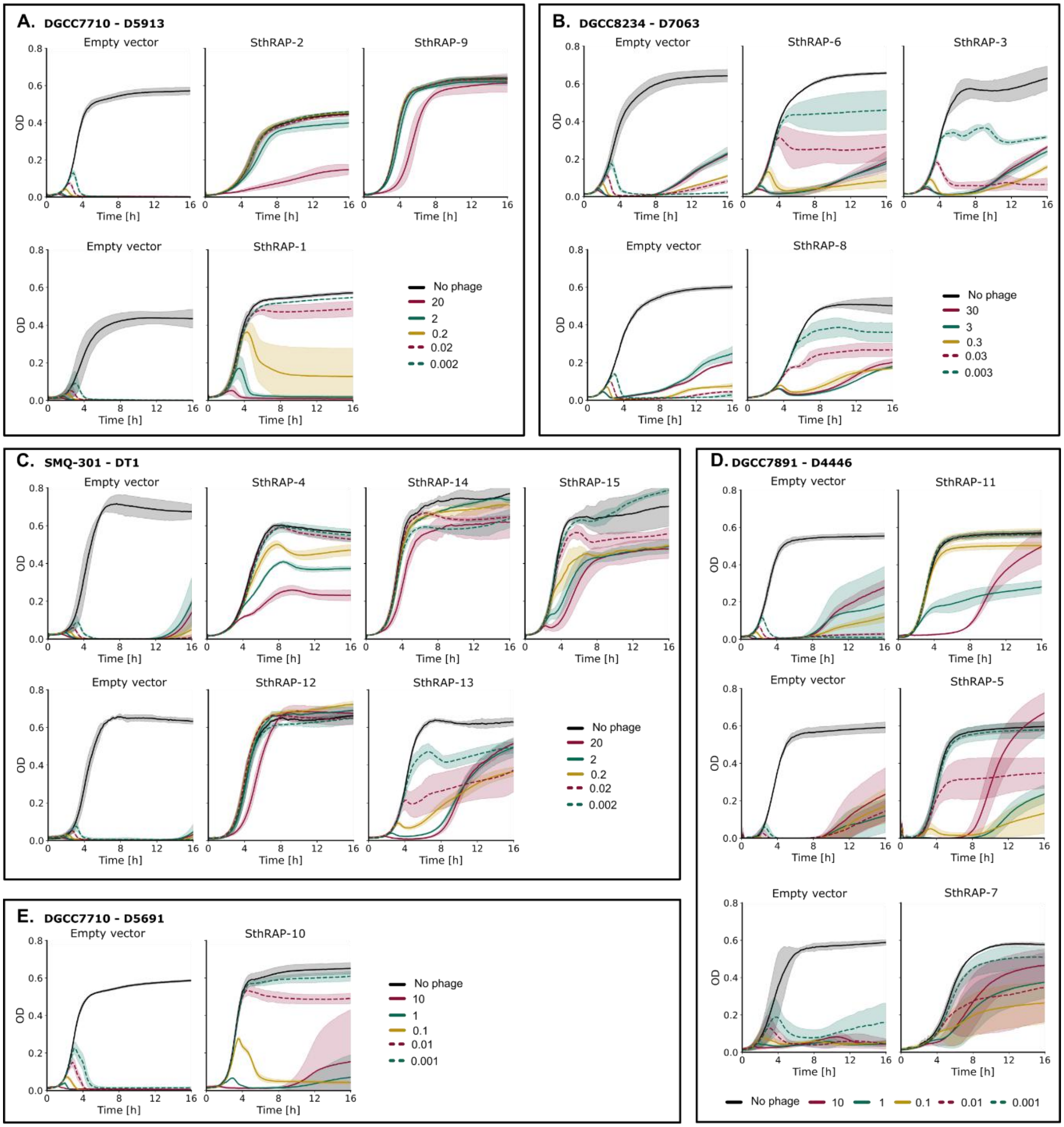


**Supplementary Figure S7 | Evaluation of anti-phage activity of SthRAP systems in liquid culture assays.** Bacterial growth was monitored over 16 hours following PHAGE infection at multiplicities of infection (MOIs) ranging from 0.001 to 30. For each defense system, the corresponding control *S. thermophilus* strain carrying the empty plasmid is displayed on the same row for comparison. Growth curves represent the mean of three biological replicates, with shaded areas indicating 95% confidence intervals. Tested bacterial strain–phage pairs include DGCC7710–D5913 (**A**), DGCC8234–D7063 (**B**), SMQ-301–DT1 (**C**), DGCC7891–D4446 (**D**), and DGCC7710–D5691 (**E**).

Internal

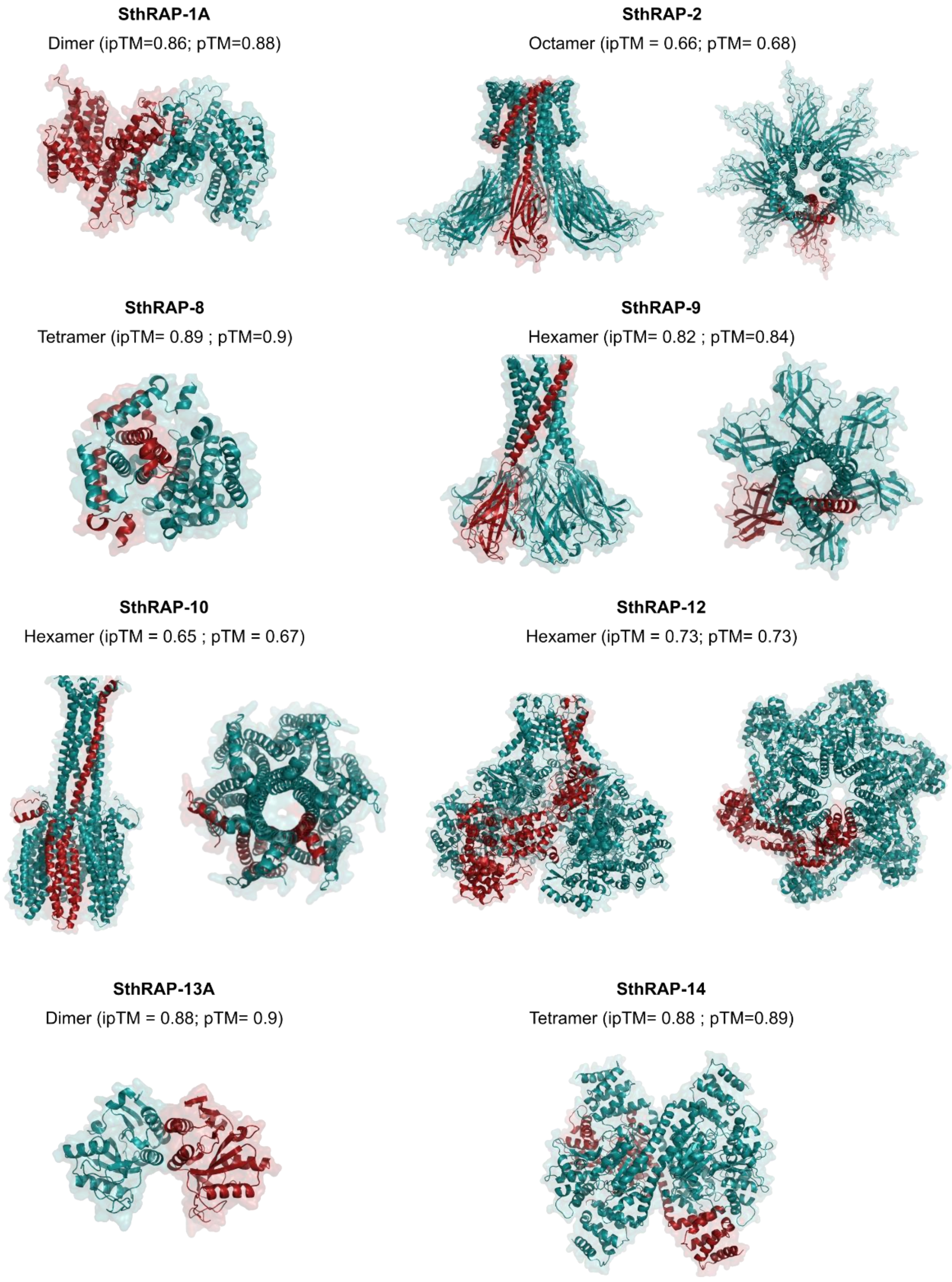


**Supplementary Figure S8 | Structural models of SthRAP system proteins predicted to form multimers by AlphaFold3.** For each system, a single monomer is highlighted in red within the predicted multimer. Structural confidence metrics are indicated as pTM (Predicted Template Modeling score) and ipTM (Interface Predicted TM score).

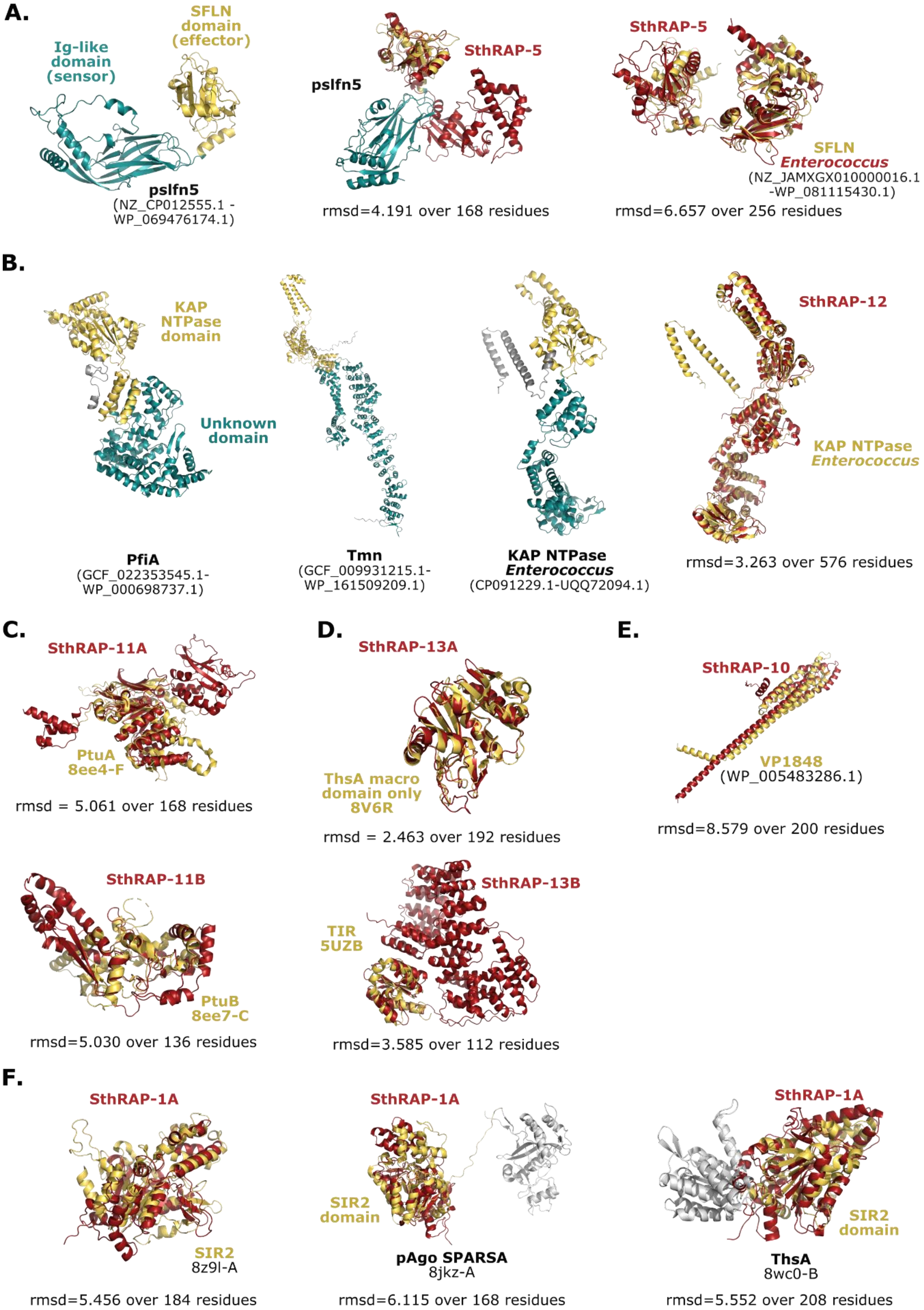
A.
SFLN
domain
(effector)
Ig-like
domain
(sensor)
pslfn5
(NZ_CP012555.1 -
WP_069476174.1)
StRAP-5
pslfn5
rmsd=4.191 over 168 residues
SthRAP-5
SFLN
Enterococcus
(NZ_JAMXGX010000016.1
-WP_081115430.1)
rmsd=6.657 over 256 residues
B.
KAP
NTPase
domain
Unknown
domain
PfiA
(GCF_022353545.1-
WP_000698737.1)
Tmn
(GCF_009931215.1-
WP_161509209.1)
KAP NTPase
Enterococcus
(CP091229.1-UQQ72094.1)
SthRAP-12
KAP NTPase
Enterococcus
rmsd=3.263 over 576 residues
C.
SthRAP-11A
PtuA
8ee4-F
rmsd = 5.061 over 168 residues
SthRAP-11B
PtuB
8ee7-C
rmsd=5.030 over 136 residues
D.
SthRAP-13A
ThsA macro
domain only
8V6R
rmsd = 2.463 over 192 residues
SthRAP-13B
TIR
5UZB
rmsd=3.585 over 112 residues
E.
SthRAP-10
VP1848
(WP_005483286.1)
rmsd=8.579 over 200 residues
F.
SthRAP-1A
SIR2
8z9l-A
rmsd=5.456 over 184 residues
SthRAP-1A
SIR2
domain
pAgo SPARSA
8jkz-A
rmsd=6.115 over 168 residues
SthRAP-1A
SIR2
domain
ThsA
8wc0-B
rmsd=5.552 over 208 residues

**Supplementary Figure S9 | Structural superimposition of SthRAP systems with known defense systems or domains found in other antiphage systems.** For proteins with experimentally determined structures, the corresponding PDB accession number is indicated. AlphaFold3 models were used for other proteins, with the strain and accession number provided in brackets. Superimpositions were performed in PyMOL using the cealign command. **A.** Comparison of SthRAP-5 with Schlafen (SFLN) proteins identified as defense systems: pslfn5[2] and the *Enterococcus* SFLN protein[3]. For pslfn5, the effector domain (SFLN) and sensor domain (Ig-like) are highlighted in yellow and blue respectively. **B.** Defense systems containing transmembrane KAP-NTPase, with the KAP-NTPase domain highlighted in yellow and the associated unknown domain in blue. PfiA is part of a two-gene system named Pfi[4]. **C.** Superimposition of SthRAP-11 proteins with the Septu subunits[5]. **D.** Superimposition of SthRAP-13A with ThsA macro domain from Thoeris[6] and SthRAP-13B with a TIR domain[7]. **E.** Superimposition of SthRAP-10 with VP1848 from *Vibrio parahaemolyticus*[8]. **F.** Alignment of SthRAP-1cdsA with SIR2 domains including those from the defense systems pAgo SPARSA and ThsA (Thoeris), with SIR2 domains highlighted in yellow and remaining regions in gray.

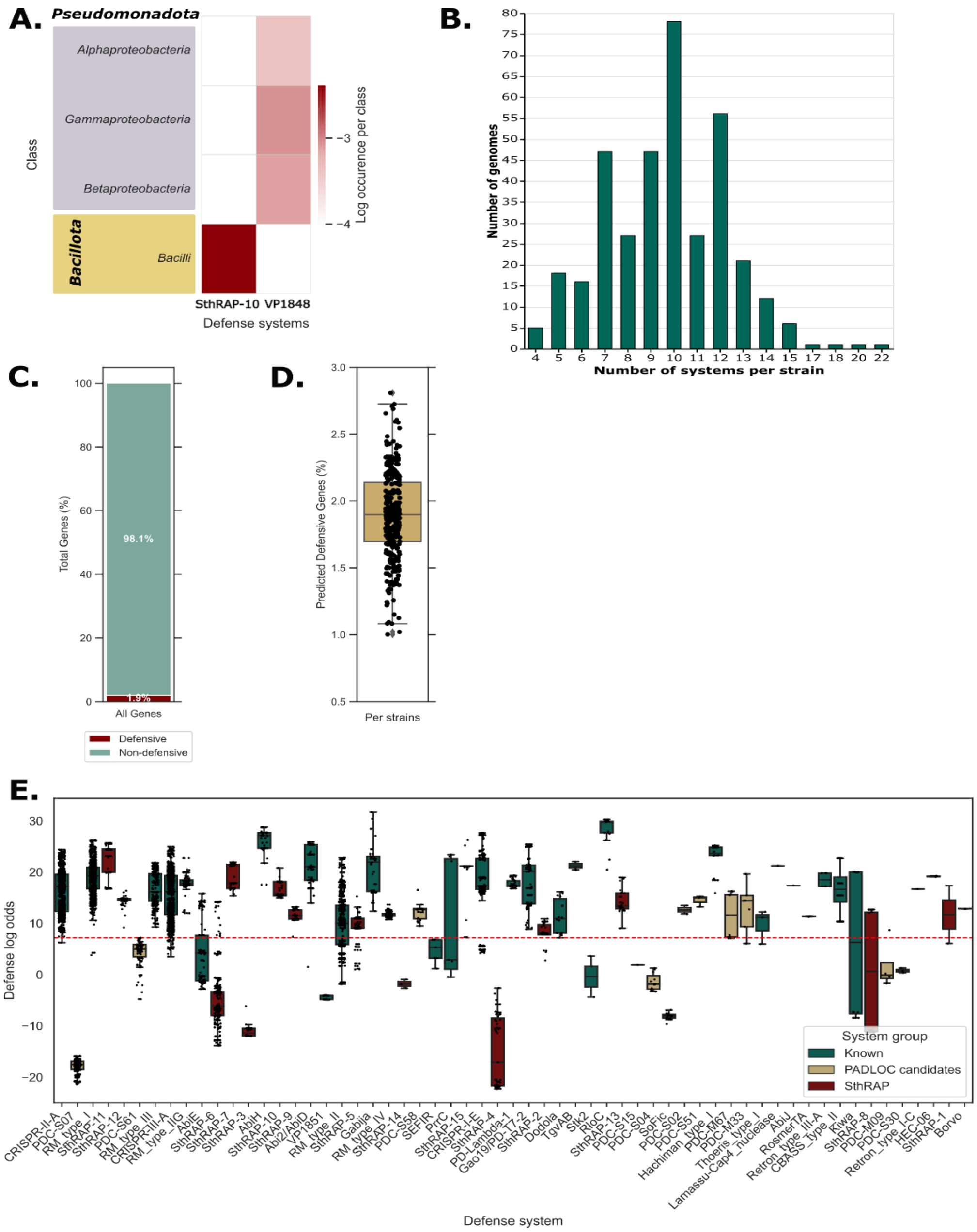


**Supplementary Figure S10 | Overall distribution of SthRAP-10 and VP1848 and prevalence analysis of predicted defense genes in *S. thermophilus* genomes. A.** Distribution of SthRAP-10 and VP1848 systems in complete bacterial genomes (RefSeq June 2025, N=50,676). These defense systems exhibit similar structure as predicted by AlphaFold3. Occurrence in each taxon is normalized to the total number of genomes in that taxon and expressed as $\log_{10}$ values. **B.** Number of validated defense systems (DefenseFinder and PADLOC systems and SthRAP systems) per genomes in *S. thermophilus* (N = 364, RefSeq June 2025). **C-D.** The same dataset was analyzed using DefensePredictor to identify predicted defense genes (log-odds > 7.2). The proportion of predicted defense genes across the entire dataset (total of 586,473 genes) (**C**) and within each individual genome (**D**) were plotted. **E.** Defense log odds score attributed by DefensePredictor to known defense genes, candidate defense genes identified by PADLOC, and SthRAP genes. Only genes superior to 7.2 were considered as defensive.

**References**


1. Gilchrist, C. L. M. & Chooi, Y. H. Clinker & clustermap. js: Automatic generation of gene cluster comparison figures. *Bioinformatics* **37,** 2473-2475 (2021).
2. Taboada, V. P. *et al.* Bacterial Schlafens mediate anti-phage defense. *bioRxiv* (2025) doi:10.1101/2025.07.24.666596.
3. Andersen, S. E. *et al.* Serine recombinases are conserved genetic markers of antiphage defense systems. *bioRxiv* (2025) doi:10.1101/2025.10.07.681051.
4. Aravind, L., Iyer, L. M., Leipe, D. D. & Koonin, E. V. A novel family of P-loop NTPases with an unusual phyletic distribution and transmembrane segments inserted within the NTPase domain. *Genome Biol.* **5**, R30 (2004).
5. Li, Y. *et al.* PtuA and PtuB assemble into an inflammasome-like oligomer for anti-phage defense. *Nat. Struct. Mol. Biol.* **31**, 413–423 (2024).
6. Shi, Y. *et al.* Structural characterization of macro domain-containing Thoeris antiphage defense systems. *Sci. Adv.* **10**, eadn3310 (2024).
7. Ve, T. *et al.* Structural basis of TIR-domain-assembly formation in MAL- and MyD88-dependent TLR4 signaling. *Nat. Struct. Mol. Biol.* **24**, 743–751 (2017).
8. Getz, L. J., Fairburn, S. R., Vivian Liu, Y., Qian, A. L. & Maxwell, K. L. Integrons are anti-phage defence libraries in *Vibrio parahaemolyticus*. *Nat. Microbiol.* **10**, 724–733 (2025).
